# Clenbuterol Attenuates Immune Reaction to Lipopolysaccharide and Its Relationship to Anhedonia in Adolescents

**DOI:** 10.1101/2021.08.07.455522

**Authors:** Tram N. B. Nguyen, Benjamin A. Ely, Danielle Pick, Manishkumar Patel, Hui Xie, Seunghee Kim-Schulze, Vilma Gabbay

## Abstract

While inflammation has been implicated in psychopathology, relationships between immune-suppressing processes and psychiatric constructs remain elusive. This study sought to assess whether β_2_-agonist clenbuterol (CBL) would attenuate immune activation in adolescents with mood and anxiety symptoms following *ex vivo* exposure of whole blood to lipopolysaccharide (LPS). Our focus on adolescents aimed to target a critical developmental period when psychiatric conditions often emerge and prior to chronicity effects. To capture a diverse range of immunologic and symptomatologic phenotypes, we included 97 psychotropic-medication free adolescents with mood and anxiety symptoms and 33 healthy controls. All participants had comprehensive evaluations and dimensional assessments of psychiatric symptoms. Fasting whole-blood samples were collected and stimulated with LPS in the presence and absence of CBL for 6 hours, then analyzed for 41 cytokines, chemokines, and hematopoietic growth factors. Comparison analyses used Bonferroni-corrected nonparametric tests. Levels of nine immune biomarkers—including IL-1RA, IL-1β, IL-6, IP-10, MCP-1, MIP-1α, MIP-1β, TGF-α, and TNF-α—were significantly reduced by CBL treatment compared to LPS alone. Exploratory factor analysis reduced 41 analytes into 5 immune factors in each experimental condition, and their relationships with psychiatric symptoms were examined as a secondary aim. CBL+LPS Factor 4—comprising EGF, PDGF-AA, PDGF-AB/BB, sCD40L, and GRO—significantly correlated with anticipatory and consummatory anhedonia, even after controlling for depression severity. This study supports the possible inhibitory effect of CBL on immune activation. Using a data-driven method, distinctive relationships between CBL-affected immune biomarkers and dimensional anhedonia were reported, further elucidating the role of β_2_-agonism in adolescent affective symptomatology.

## 1. Introduction

Inflammation has been implicated in numerous neuropsychiatric conditions [1, 2]. Work by our group and others have consistently reported a link between peripheral inflammatory processes across psychiatric conditions and ages [3–11]. The relationships between inflammation and psychopathologies are thought to reflect similar neurobiological mechanisms to those underlying “sickness behavior” in model animals [8]. Sickness behavior is characterized by fatigue, malaise, and reductions in appetite, ability to concentrate, mood, social interactions, and pleasure-seeking behaviors [12–15]. While sickness behavior is an adaptive process involving complex brain-body interactions, dysregulated immune activation is postulated to induce maladaptive changes in the central nervous system, which ultimately manifest as psychiatric disturbances [8]. Since psychiatric conditions are heterogenous, our group and others have utilized the Research Domain Criteria (RDoC) framework, which focuses on behavioral constructs rather than categorical psychiatric diagnoses [16]. We found that only anhedonia—the decreased capacity to experience pleasure—was associated with peripheral inflammation among adolescents [5, 6, 17]. As anhedonia clinically reflects reward dysfunction, our findings suggest that inflammation may induce alterations in neurocircuits underlying reward function. Indeed, our neuroimaging work supported this investigative direction, providing evidence for the relationships between clusters of immune analytes and alterations of the neural function across reward processes [18–20].

As inflammation involves many intertwined and tightly orchestrated systems, it is crucial to examine processes modulating immune activity to delineate the brain-immune relationships. One prominent immune regulator is the adrenergic nervous system, which has been shown to halt overactivation of peripheral and central immune cells through β_2_-adrenergic receptor transduction pathways [21]. Attempts to study the role of adrenergic regulation in depressive pathophysiology have yielded promising results. Earlier post-mortem studies showed differential adrenoreceptor density in reward-related cortical regions between antidepressant-free suicide victims and matched controls [22, 23]. More recently, catecholamine depletion has been linked to alterations in reward processing among adults with major depressive disorder [24]. Nevertheless, the anti-inflammatory effects of the adrenergic system on reward function remains largely unexamined in depression and across the lifespan. Among immune-inhibitory β_2_ receptor agonists, clenbuterol (CBL) is a promising candidate for use in modeling adrenergic regulation of immune activation *ex vivo*. Especially relevant to psychiatric contexts, CBL readily crosses the blood brain barrier to act on central β_2_ receptors, with preclinical evidence indicating that CBL reduces neuroinflammation associated with strokes [25] and exhibits antidepressant-like effects [26] in rodent models. However, investigation of β_2_ agonists adrenergic system regulating immune CBL’s anti-inflammatory property using human blood samples is scarce, and available studies have only analyzed a limited number of inflammatory biomarkers.

Building upon the above observations, here we sought to investigate whether adrenergic immune modulation could be achieved in a functional immune response model using lipopolysaccharide (LPS) and CBL in whole blood. In alignment with the RDoC framework, we examined behavioral constructs to capture the full range of symptom severity rather than stratifying participants by categorical diagnoses. Therefore, participants were psychotropic-medication free adolescents with diverse mood and anxiety symptoms as well as healthy adolescents. Our focus on adolescence aimed to capture a critical period with increased vulnerability to psychopathology development. We used a comprehensive profile of 41 immune biomarkers including cytokines, chemokines, and hematopoietic growth factors. We hypothesized that CBL would attenuate immune activation following exposure to LPS in adolescents. There were no *a priori* hypotheses regarding which specific LPS-stimulated immune proteins would be downregulated by CBL, as we expected a general attenuating effect on immune protein production. Further, based on our prior finding implicating inflammation in anhedonia, we also conducted secondary analyses assessing the relationships between CBL-induced attenuation of inflammatory effects and severity of depression, anxiety, and anhedonia symptoms.

## 2. Methods

### 2.1. Study Participants

Adolescents in the New York City area were recruited via community advertisements and clinician referrals. On the first visit, adolescents underwent a comprehensive medical and psychiatric assessment to determine eligibility for study participation. Eligible adolescents were instructed to return to the study site within approximately 2 weeks to undergo study procedures, including vital sign measurements, a fasting blood draw, a urine toxicology screen, a pregnancy test if female, and completion of clinical questionnaires.

This study was approved by the Institutional Review Boards at Icahn School of Medicine at Mount Sinai and Albert Einstein College of Medicine. Participants ages 18 and older (n=24) provided written informed consent; participants younger than 18 years old provided assent, and a legal guardian provided signed consent.

### 2.2. Inclusion and Exclusion Criteria

The study included medically healthy adolescents ages 12 to 20 years. Inclusion criteria for the clinical group: primary presentation of mood and anxiety symptoms regardless of meeting diagnostic criteria per the Diagnostic and Statistical Manual of Mental Disorders (DSM), Fourth or Fifth Edition [27, 28]. Exclusion criteria for the clinical group: a) current or past DSM diagnosis of eating disorders, schizophrenia, pervasive developmental disorder, or substance use disorder, b) intake of any psychotropic medication within 30-90 days (depending on drug half-life) prior to the blood draw; c) history of immunological or hematological disorder; d) intake of any immune-affecting medication and supplements (e.g., steroids, non-steroidal anti-inflammatory drugs, omega-3 fatty acids) within 2 weeks prior to study enrollment; c) history of chronic fatigue syndrome; d) any infection during the month prior to the blood draw (including the common cold); e) history of significant medical or neurological disorders; f) an estimated IQ below 80 based on the Kaufmann Brief Intelligence Test [KBIT; 29]; g) positive urine toxicology screens; h) in females, a positive pregnancy test.

Healthy controls (HC) were required to meet all above exclusion criteria in addition to be without a lifetime history of any psychiatric diagnoses, including subthreshold symptoms as determined by clinical evaluations (detailed in section *2.3*. below).

### 2.3. Clinical Evaluations

Each participant received a thorough evaluation consisting of a comprehensive psychiatric evaluation, medical history as well as laboratory tests including complete blood count, metabolic panel, liver and thyroid function tests, and a urine toxicology test. Both the participating adolescent and their accompanying parent were interviewed by a clinician to assess for possible presence of an infectious illness (including the common cold) within the month immediately prior to enrollment.

Psychiatric diagnostic evaluation: The semi-structured Schedule for Affective Disorders and Schizophrenia – Present and Lifetime version [K-SADS-PL; 30], an interview-based assessment tool commonly used in pediatric research settings, was conducted by a trained, licensed psychiatrist or clinical psychologist to assess presence of any psychiatric symptoms and lifetime history of psychiatric diagnoses per the DSM. Evaluations were discussed between the interviewing clinician and the Principal Investigator, a board-certified child and adolescent psychiatrist, to enhance diagnostic reliability. The K-SADS-PL was utilized to ensure participants meet study eligibility and determine if they belonged to the clinical or HC group.

Dimensional symptom assessment: The self-rated, 39-item Multidimensional Anxiety Scale for Children [MASC; 31] was employed to quantify anxiety severity (scores ranging from 0 to 117). Participants self-reported their depression severity with the 21-item Beck Depression Inventory, 2^nd^ edition [BDI-II; scores ranging 0 to 63; 32]. Higher scores on the MASC and BDI indicate more severe anxiety and depression, respectively. Additionally, we utilized the Temporal Experience of Pleasure Scale [TEPS; 33] to separately capture anticipatory (TEPS-A; scores ranging 10 to 60) and consummatory (TEPS-C; scores ranging 8 to 48) components of anhedonia. Higher TEPS scores suggest lower levels of anhedonia. The TEPS’ internal consistency is comparable in both adult and adolescent samples [33, 34], and its breakdown of anticipatory and consummatory subscales has been supported by exploratory factor analysis using data collected in two independent samples [35]. We note that the BDI, MASC, and TEPS are designed to dimensionally probe the full range of symptoms severity, and thus HCs in our sample were expected to report non-zero, though non-pathological, levels of depression, anxiety, and anhedonia (see **Table 1**).

**Table 1.**
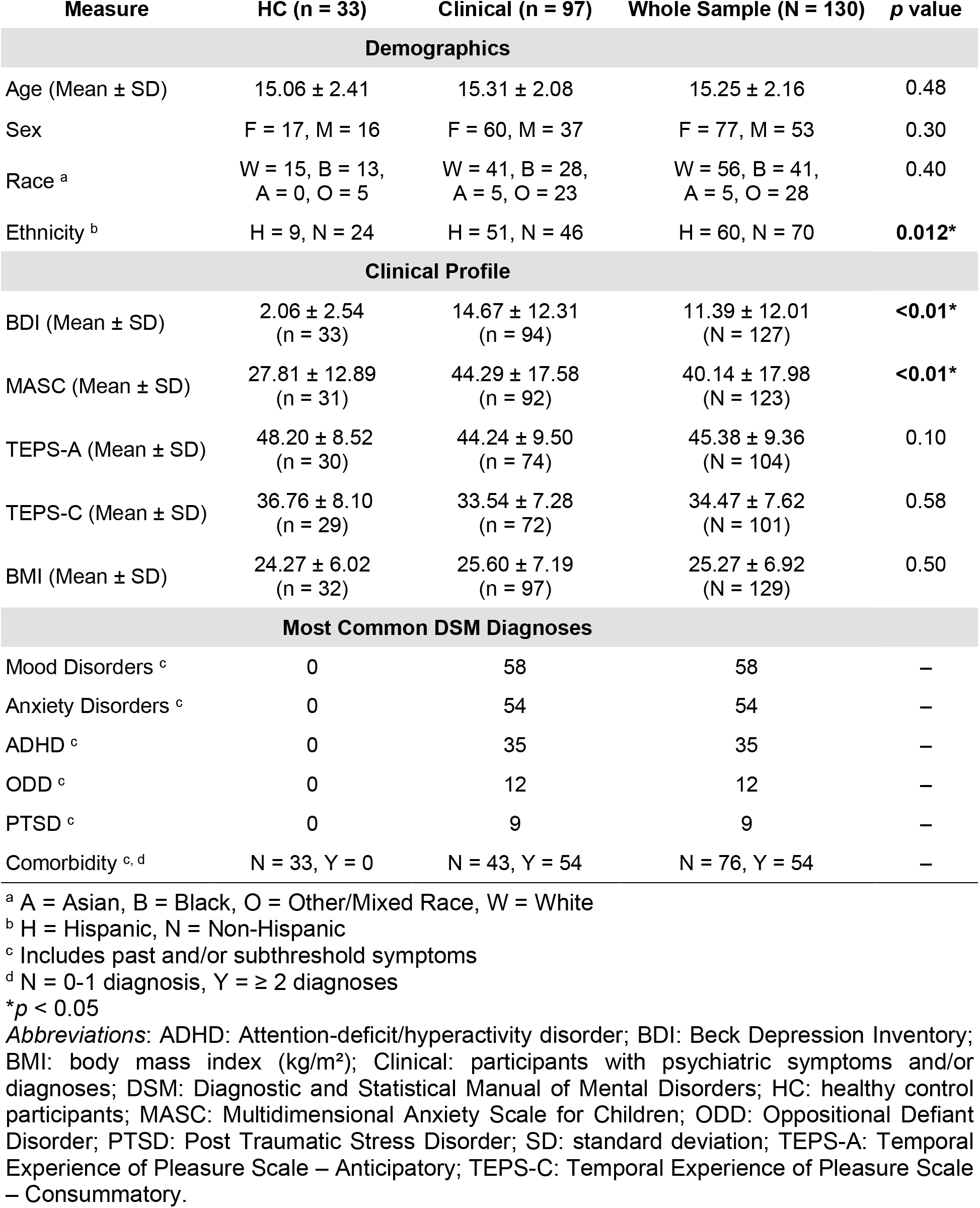
Demographic and clinical characteristics of study participants. Reported *p* values are of appropriate bivariate tests comparing the clinical and healthy control groups.

| Measure | HC (n = 33) | Clinical (n = 97) | Whole Sample (N = 130) | p value |
| --- | --- | --- | --- | --- |
| **Demographics** |  |  |  |  |
| Age (Mean ± SD) | 15.06 ± 2.41 | 15.31 ± 2.08 | 15.25 ± 2.16 | 0.48 |
| Sex | F = 17, M = 16 | F = 60, M = 37 | F = 77, M = 53 | 0.30 |
| Race <sup>a</sup> | W = 15, B = 13,<br>A = 0, O = 5 | W = 41, B = 28,<br>A = 5, O = 23 | W = 56, B = 41,<br>A = 5, O = 28 | 0.40 |
| Ethnicity <sup>b</sup> | H = 9, N = 24 | H = 51, N = 46 | H = 60, N = 70 | **0.012\*** |
| **Clinical Profile** |  |  |  |  |
| BDI (Mean ± SD) | 2.06 ± 2.54<br>(n = 33) | 14.67 ± 12.31<br>(n = 94) | 11.39 ± 12.01<br>(N = 127) | **<0.01\*** |
| MASC (Mean ± SD) | 27.81 ± 12.89<br>(n = 31) | 44.29 ± 17.58<br>(n = 92) | 40.14 ± 17.98<br>(N = 123) | **<0.01\*** |
| TEPS-A (Mean ± SD) | 48.20 ± 8.52<br>(n = 30) | 44.24 ± 9.50<br>(n = 74) | 45.38 ± 9.36<br>(N = 104) | 0.10 |
| TEPS-C (Mean ± SD) | 36.76 ± 8.10<br>(n = 29) | 33.54 ± 7.28<br>(n = 72) | 34.47 ± 7.62<br>(N = 101) | 0.58 |
| BMI (Mean ± SD) | 24.27 ± 6.02<br>(n = 32) | 25.60 ± 7.19<br>(n = 97) | 25.27 ± 6.92<br>(N = 129) | 0.50 |
| **Most Common DSM Diagnoses** |  |  |  |  |
| Mood Disorders <sup>c</sup> | 0 | 58 | 58 | – |
| Anxiety Disorders <sup>c</sup> | 0 | 54 | 54 | – |
| ADHD <sup>c</sup> | 0 | 35 | 35 | – |
| ODD <sup>c</sup> | 0 | 12 | 12 | – |
| PTSD <sup>c</sup> | 0 | 9 | 9 | – |
| Comorbidity <sup>c, d</sup> | N = 33, Y = 0 | N = 43, Y = 54 | N = 76, Y = 54 | – |

<sup>a</sup> A = Asian, B = Black, O = Other/Mixed Race, W = White
<sup>b</sup> H = Hispanic, N = Non-Hispanic
<sup>c</sup> Includes past and/or subthreshold symptoms
<sup>d</sup> N = 0-1 diagnosis, Y = ≥ 2 diagnoses
\**p* < 0.05
**Abbreviations:** ADHD: Attention-deficit/hyperactivity disorder; BDI: Beck Depression Inventory; BMI: body mass index (kg/m<sup>2</sup>); Clinical: participants with psychiatric symptoms and/or diagnoses; DSM: Diagnostic and Statistical Manual of Mental Disorders; HC: healthy control participants; MASC: Multidimensional Anxiety Scale for Children; ODD: Oppositional Defiant Disorder; PTSD: Post Traumatic Stress Disorder; SD: standard deviation; TEPS-A: Temporal Experience of Pleasure Scale – Anticipatory; TEPS-C: Temporal Experience of Pleasure Scale – Consummatory.

### 2.4. *Ex Vivo* Immune Stimulation with Lipopolysaccharide and Clenbuterol

Blood samples were collected between 9 and 10 A.M. after an overnight fast lasting at least 12 hours. Functional immune responses were assessed *in vitro* using a well-established LPS challenge protocol that reliably induces pro-inflammatory cytokine production [36]. Whole blood samples were cultured on RPMI Medium 1640 (1×) with L-Glutamine, Cat# 11875-093 (Invitrogen, Waltham, MA) for 6 hours at 37°C in a 5% CO_2_ tissue culture incubator under three conditions: in the presence of LPS alone, in the presence of LPS and the β_2_-agonist CBL, and in the absence of both LPS and CBL (i.e. Control condition). Specifically, LPS-EB Ultrapure 5mg, 5 × 10^6^ EU/ml, Cat# tlrl-pelps (InvivoGen, San Diego, CA) with working concentration of 0.1 μg/mL and Clenbuterol hydrochloride, Cat# C5423-10MG (MilliporeSigma, St. Louis, MO) with working concentration of 10^−6^ M were used. After 6 hours of incubation, the cultured whole blood samples were centrifuged. Supernatant was collected and stored at −20°C. Comprehensive details on sample collection, storage, dose determination of LPS and CBL, as well as experimental environment can be found in **Supplementary Methods** – *SM1*. Additionally, details of secondary analyses to determine if storage time affected immune protein levels within our sample were reported in **Supplementary Methods** – *SM2*.

Levels of 41 immune analytes (cytokines, chemokines, and growth factors; see **Supplementary Table 1**) were measured using a bead-based Luminex-200 system and the XMap Platform (Luminex Corporation, Austin, TX), as described in **Supplementary Methods** – *SM2*. Assays were performed at the Mount Sinai Human Immune Monitoring Center by laboratory staff blinded to participants’ clinical status. The combination of assays has been previously validated using secondary measurements [37–41]. Duplicates of median fluorescent intensity (MFI) values were measured for each of the 3 peripheral whole blood culture growth conditions (Control, LPS, LPS+CBL), and mean MFI values were generated. While absolute concentration values (**Supplementary Table 1**) were computed from mean MFI values, we opted to use MFI values in our analyses as these allow for increased statistical power [42, 43]. To enhance the reproducibility and transparency of our findings, we reported further details on the detection limits, methods to account for potential assay drifts, and reasons for missing MFI values for each analyte in **Supplementary Methods** – *SM2* and *SM3*.

### 2.5. Statistical Analyses

Primary analyses were conducted in MATLAB R2020b (MathWorks, Natick, MA). An overview of our analytic protocol can be found in **Supplementary Figure 2**. Shapiro-Wilk tests indicated a non-normal distribution for most analytes in our sample, necessitating a nonparametric approach. We first utilized Friedman tests, controlling for family-wise error rate using Bonferroni correction, to test for significant differences in analyte levels across the three conditions (Control, LPS, and LPS+CBL). For any analyte found to differ significantly, *post-hoc* Wilcoxon signed-rank tests with Bonferroni correction were used to examine pairwise differences between conditions. Effect sizes for Wilcoxon signed-rank tests were computed as the correlation coefficient 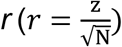 [44–47].

In secondary analyses, given the unknown relations between CBL-attenuated immune biomarkers and subjective psychiatric symptoms, we utilized an exploratory factor analysis (EFA) model to identify sources of shared variance within subjects’ detailed immune profiles in a data-driven manner. Immune analyte data were examined using 3 discrete EFA models corresponding to the 3 conditions (Control, LPS, LPS+CBL). EFA was performed in SPSS 27.0 (IBM, Armonk, NY) using principal component analysis (PCA) dimensionality reduction and varimax orthogonal rotation. Finally, Spearman partial correlations controlling for age, sex, and body mass index (BMI) were used to assess the relationships between each immune factor scores and BDI (depression), MASC (anxiety), TEPS-A (anticipatory anhedonia), and TEPS-C (consummatory anhedonia) scores. Spearman correlations between immune factors and anhedonia measures were repeated while controlling for depression severity.

Following our analyses outlined above, we performed a supplementary factor analysis to assess relative changes in analyte levels between LPS and LPS+CBL conditions and related the resulting factors to anhedonia scores (see **Supplementary Results** – *SR1*). Further *post hoc* analyses were conducted to contrast immune analyte relationships between LPS+CBL and LPS conditions. Spearman correlations (*rho*) between pairs of analytes in the 2 conditions were computed, *z*-transformed, subtracted from each other, then transformed back to *rho* coefficients. This yielded a total of 820 pairwise correlation differences (Δ*rho*) per subject. The significance of each Δ*rho* was assessed using a bias-corrected and accelerated bootstrap [48–50] with 10^5^ resamples. We additionally utilized this bootstrapping method to determine whether observed relationships between symptom measures and immune factors are significantly stronger than other comparisons (details in **Supplementary Results** – *SR2*).

## 3. Results

We reported characteristics of our sample and the results of our primary aim investigating adrenergic-induced immune responses in subsections *3.1* and *3.2*. In subsections *3.3* and *3.4*, we presented results of our secondary aim probing potential relationships between immune attenuation and affective symptomatology.

### 3.1. Participants’ Characteristics

This study enrolled a total of 130 adolescents, including 97 in the clinical group (85 endorsed mood and anxiety symptoms; 12 with other externalizing behavioral symptoms without comorbid mood and anxiety symptoms) and 33 HC. As summarized in **Table 1**, participants were racially and ethnically diverse. Demographically, participants with psychiatric symptoms and HC were similar (*p* > 0.1), except for a significantly higher proportion of Hispanic individuals among the clinical sample (*p* = 0.012). More than half of participants in the clinical group had at least two concurrent DSM diagnoses. As expected, adolescents in the clinical group exhibited higher depression and anxiety severity compared to HC (see **Table 1**). Depression and anhedonia scores were skewed, while anxiety scores were normally distributed (see **Supplementary Figure 3**). Across all participants, anticipatory and consummatory anhedonia scores were significantly correlated with each other and with depression, but not with anxiety (see **Supplementary Table 2**).

### 3.2. Lipopolysaccharide and Clenbuterol Effects on Immune Analytes

Immune analyte levels were not significantly different between HC and participants with psychiatric symptoms for any condition (see **Supplementary Table 3**). Consequently, all analyses used combined data from the whole sample. Friedman tests (detailed in **Supplementary Table 4**) indicated significant differences in 21 analytes across the three conditions after Bonferroni correction 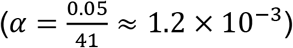.

Pairwise follow-up analyses (i.e., Control vs. LPS, LPS vs. LPS+CBL, Control vs. LPS+CBL) of these 21 analytes using Wilcoxon signed-rank tests are detailed in **Table 2**, with results considered significant at the Bonferroni corrected 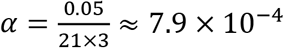 level. Relative to the Control condition, LPS increased levels of 19 out of the 21 analytes, with only PDGF-AB/BB and sCD40L not meeting significance. Levels of 17/21 analytes (all except Fractalkine, G-CSF, IL-12P40, and sCD40L) were also elevated in the LPS+CBL condition relative to Control. For all but 2/21 analytes (PDGF-AB/BB and sCD40L), effect sizes were smaller for LPS+CBL vs. Control than for LPS vs. Control, consistent with the immunosuppressant activity of CBL. Compared to LPS, levels of 9/21 analytes were significantly reduced in the LPS+CLB condition, while only 1/21 (sCD40L) was found to be increased in the presence of CBL. Analytes with the greatest differences in expression between LPS+CBL and LPS conditions included MIP-1β (*r* = −0.73), TNF-α (*r* = −0.70), MIP-1α (*r* = −0.65), and IL-6 (*r* = −0.62). These findings remained consistent when stratified by group membership (see **Supplementary Tables 5–6**).

**Table 2.** *Post hoc* pairwise comparisons of immune analyte levels across 3 *ex vivo* conditions in whole sample (N=130). ΔMFI: difference of median MFI levels across pairs of conditions

| Immune Marker | LPS vs. Control |  |  | LPS+CBL vs. LPS |  |  | LPS+CBL vs. Control |  |  |
| --- | --- | --- | --- | --- | --- | --- | --- | --- | --- |
| | ΔMFI | r | p value | ΔMFI | r | p value | ΔMFI | r | p value |
| Fractalkine | 1 | 0.40 | $3.5 \times 10^{-6} *$ | -0.25 | -0.21 | $1.7 \times 10^{-2}$ | 0.75 | 0.24 | $5.3 \times 10^{-3}$ |
| G-CSF | 3.25 | 0.34 | $9.5 \times 10^{-5} *$ | -0.75 | -0.10 | $2.3 \times 10^{-1}$ | 2.5 | 0.23 | $1.0 \times 10^{-2}$ |
| GM-CSF | 1.75 | 0.40 | $3.5 \times 10^{-6} *$ | 0.125 | -0.15 | $6.9 \times 10^{-2}$ | 1.875 | 0.30 | $6.3 \times 10^{-4} *$ |
| GRO<sup>a</sup> | 200.875 | 0.42 | $1.0 \times 10^{-6} *$ | 448.275 | 0.22 | $1.1 \times 10^{-2}$ | 649.15 | 0.42 | $6.9 \times 10^{-7} *$ |
| IL-10 | 2.25 | 0.54 | $1.2 \times 10^{-10} *$ | 1 | -0.01 | $7.7 \times 10^{-1}$ | 3.25 | 0.54 | $1.1 \times 10^{-10} *$ |
| IL-12P40 | 2.75 | 0.36 | $3.7 \times 10^{-5} *$ | -0.875 | -0.22 | $1.0 \times 10^{-2}$ | 1.875 | 0.23 | $9.5 \times 10^{-3}$ |
| IL-1RA | 23.75 | 0.73 | $5.2 \times 10^{-21} **$ | -10.5 | -0.47 | $3.7 \times 10^{-8} *$ | 13.25 | 0.57 | $8.2 \times 10^{-12} **$ |
| IL-1α | 3.5 | 0.41 | $2.2 \times 10^{-6} *$ | -1.5 | -0.21 | $1.2 \times 10^{-2}$ | 2 | 0.31 | $3.1 \times 10^{-4} *$ |
| IL-1β | 46 | 0.82 | $7.8 \times 10^{-29} **$ | -12.875 | -0.51 | $1.0 \times 10^{-9} *$ | 33.125 | 0.77 | $3.4 \times 10^{-24} **$ |
| IL-4 | 1.875 | 0.43 | $4.4 \times 10^{-7} *$ | 1 | 0.039 | $6.6 \times 10^{-1}$ | 2.875 | 0.39 | $6.8 \times 10^{-6} *$ |
| IL-6 | 136.75 | 0.82 | $8.2 \times 10^{-29} **$ | -47 | -0.62 | $2.6 \times 10^{-14} **$ | 89.75 | 0.76 | $7.0 \times 10^{-23} **$ |
| IL-7 | 2.25 | 0.42 | $1.6 \times 10^{-6} *$ | 0 | -0.12 | $1.8 \times 10^{-1}$ | 2.25 | 0.36 | $3.5 \times 10^{-5} *$ |
| IL-8 | 380 | 0.82 | $3.4 \times 10^{-28} **$ | -31.25 | -0.26 | $2.8 \times 10^{-3}$ | 348.75 | 0.79 | $3.0 \times 10^{-25} **$ |
| IP-10 | 92.375 | 0.71 | $3.5 \times 10^{-19} **$ | -66.375 | -0.56 | $2.4 \times 10^{-11} **$ | 26 | 0.35 | $4.8 \times 10^{-5} *$ |
| MCP-1 | 378.3 | 0.71 | $2.6 \times 10^{-19} **$ | -205.55 | -0.60 | $4.8 \times 10^{-13} **$ | 172.75 | 0.37 | $2.3 \times 10^{-5} *$ |
| MIP-1α | 201 | 0.77 | $3.7 \times 10^{-23} **$ | -138.75 | -0.65 | $8.3 \times 10^{-16} **$ | 62.25 | 0.64 | $3.0 \times 10^{-15} **$ |
| MIP-1β | 322.375 | 0.82 | $1.2 \times 10^{-28} **$ | -229.375 | -0.73 | $2.4 \times 10^{-20} **$ | 93 | 0.72 | $8.7 \times 10^{-20} **$ |
| PDGF-AB/BB<sup>a</sup> | 23 | 0.23 | $8.0 \times 10^{-3}$ | 82.25 | 0.21 | $1.5 \times 10^{-2}$ | 105.25 | 0.33 | $1.8 \times 10^{-4} *$ |
| sCD40L<sup>a</sup> | 21.625 | 0.21 | $1.5 \times 10^{-2}$ | 42.75 | 0.33 | $2.0 \times 10^{-4} *$ | 64.375 | 0.28 | $1.4 \times 10^{-3}$ |
| TGF-α | 4.5 | 0.65 | $8.6 \times 10^{-16} **$ | 0.25 | -0.36 | $2.7 \times 10^{-5} *$ | 4.75 | 0.54 | $1.1 \times 10^{-10} *$ |
| TNF-α | 123.5 | 0.84 | $8.6 \times 10^{-32} **$ | -71.375 | -0.70 | $1.0 \times 10^{-18} **$ | 52.125 | 0.78 | $1.7 \times 10^{-24} **$ |
| EGF<sup>a</sup> | 26.625 | — | — | 10.875 | — | — | 37.5 | — | — |
| PDGF-AA<sup>a</sup> | 199.1 | — | — | 132.15 | — | — | 331.25 | — | — |

### 3.3. Immune Analyte Dimensionality Reduction Across Experimental Conditions

In each of the 3 examined conditions (see **Methods** – *2.5*.), EFA yielded an initial model containing 8 orthogonal immune factors with eigenvalues greater than 1, of which 5 factors were retained for our analyses on the basis of variance explained (≥5%) and scree plots. The factors explained 81.1%, 73.5%, and 74.1% variance of the original immune analyte data for the Control, LPS, and LPS+CBL conditions, respectively. Across all conditions, similar immune analytes consistently loaded on the same factors. Factor loading scores can be found in **Table 3**. For ease of interpretation, factors in the LPS and LPS+CBL conditions were listed using the same numbering scheme as that of Control factors throughout the study.

**Table 3.** Factor loadings for 5 factors (F1 – F5) derived from cytokine levels in the control, LPS, and LPS + CBL conditions using principal component analysis (Rotation method: Varimax with Kaiser Normalization). These 5 factors explained 81.1%, 73.5%, and 74.1% data variance, respectively. Major loadings (>0.3) for each factor are bolded. *For ease of interpretation, all factors in LPS and LPS+CBL conditions were listed following the same numbering scheme as Control factors.

| Cytokine | Control |  |  |  |  | LPS |  |  |  |  | LPS + CBL |  |  |  |  |
| --- | --- | --- | --- | --- | --- | --- | --- | --- | --- | --- | --- | --- | --- | --- | --- |
|  | F1 | F2 | F3 | F4 | F5 | F1 | F2 | F3 | F4 | F5 | F1 | F2 | F3 | F4 | F5 |
| IL-3 | **0.956** | 0.071 | 0.021 | -0.004 | 0.047 | **0.871** | 0.084 | 0.093 | 0.084 | 0.029 | **0.882** | 0.101 | 0.082 | 0.108 | 0.002 |
| IL-12P70 | **0.945** | 0.193 | 0.104 | -0.049 | 0.056 | **0.778** | 0.466 | 0.071 | 0.061 | -0.004 | **0.754** | 0.447 | 0.203 | 0.108 | 0.041 |
| Flt-3L | **0.943** | 0.083 | 0.092 | -0.012 | 0.058 | **0.737** | 0.277 | 0.035 | 0.144 | -0.021 | **0.663** | 0.312 | 0.179 | 0.214 | 0.016 |
| FGF-2 | **0.942** | 0.137 | 0.095 | 0.077 | 0.045 | **0.747** | 0.401 | 0.013 | 0.285 | -0.032 | **0.710** | 0.366 | 0.142 | 0.372 | 0.015 |
| IFN-α2 | **0.919** | 0.192 | 0.152 | 0.061 | 0.103 | **0.735** | 0.367 | 0.087 | 0.234 | 0.076 | **0.704** | 0.330 | 0.130 | 0.275 | 0.146 |
| IL-7 | **0.916** | 0.189 | 0.135 | 0.165 | 0.111 | **0.653** | 0.425 | 0.192 | 0.370 | 0.093 | **0.537** | 0.520 | 0.134 | 0.333 | 0.148 |
| VEGF | **0.913** | 0.332 | 0.124 | 0.070 | 0.093 | **0.761** | 0.537 | 0.087 | 0.164 | 0.093 | **0.724** | 0.540 | 0.163 | 0.220 | 0.113 |
| IL-15 | **0.877** | 0.277 | -0.024 | -0.068 | 0.150 | **0.687** | 0.364 | -0.022 | -0.028 | 0.245 | **0.697** | 0.411 | -0.106 | -0.017 | 0.191 |
| IL-12P40 | **0.870** | 0.306 | 0.043 | -0.097 | 0.146 | **0.764** | 0.297 | 0.101 | -0.127 | 0.204 | **0.747** | 0.386 | -0.041 | -0.063 | 0.178 |
| IL-2 | **0.863** | 0.249 | 0.039 | 0.098 | -0.005 | **0.859** | 0.139 | -0.024 | 0.099 | 0.013 | **0.882** | 0.122 | 0.073 | 0.143 | 0.004 |
| GM-CSF | **0.856** | 0.353 | 0.264 | 0.020 | 0.117 | 0.562 | **0.718** | 0.202 | 0.082 | 0.051 | 0.541 | **0.729** | 0.179 | 0.129 | 0.094 |
| G-CSF | **0.825** | 0.097 | 0.070 | 0.317 | 0.133 | -0.047 | 0.106 | -0.022 | **0.269** | 0.028 | **0.551** | 0.513 | 0.119 | 0.469 | 0.099 |
| IFN-γ | **0.820** | 0.185 | 0.222 | 0.006 | 0.051 | **0.734** | 0.247 | 0.182 | 0.017 | 0.029 | **0.734** | 0.223 | 0.295 | 0.062 | 0.047 |
| IL-10 | **0.809** | 0.173 | 0.046 | -0.093 | 0.093 | **0.622** | 0.183 | 0.366 | -0.175 | 0.163 | **0.625** | 0.316 | 0.252 | -0.109 | 0.092 |
| IL-17A | **0.762** | 0.399 | 0.120 | -0.038 | -0.025 | **0.621** | 0.531 | 0.116 | -0.005 | -0.058 | **0.648** | 0.425 | 0.254 | 0.045 | -0.036 |
| Fractalkine | **0.694** | 0.152 | 0.049 | 0.163 | 0.045 | **0.476** | 0.230 | 0.015 | 0.300 | -0.004 | **0.391** | 0.197 | 0.050 | 0.333 | 0.200 |
| MDC | **0.202** | 0.055 | 0.049 | 0.100 | -0.053 | 0.186 | **0.644** | -0.008 | 0.079 | -0.098 | 0.103 | **0.474** | -0.071 | 0.056 | -0.086 |
| MCP-3 | 0.269 | **0.938** | -0.011 | 0.025 | 0.100 | 0.426 | **0.816** | -0.032 | 0.053 | 0.221 | 0.483 | **0.794** | -0.083 | 0.016 | 0.211 |
| IL-13 | 0.293 | **0.930** | -0.033 | 0.048 | 0.172 | 0.419 | **0.797** | -0.097 | 0.080 | 0.337 | 0.486 | **0.756** | -0.121 | 0.036 | 0.317 |
| TNF-β | 0.249 | **0.912** | -0.035 | 0.036 | 0.294 | 0.392 | **0.759** | -0.110 | 0.085 | 0.442 | 0.456 | **0.717** | -0.125 | 0.031 | 0.439 |
| IL-1RA | 0.311 | **0.910** | 0.056 | 0.067 | 0.153 | 0.489 | **0.722** | 0.082 | 0.052 | 0.257 | 0.540 | **0.709** | 0.000 | 0.012 | 0.261 |
| IL-5 | 0.315 | **0.891** | -0.053 | 0.072 | 0.145 | 0.433 | **0.748** | -0.104 | 0.113 | 0.299 | 0.496 | **0.697** | -0.138 | 0.051 | 0.287 |
| TGF-α | 0.467 | **0.761** | 0.089 | -0.051 | -0.012 | 0.424 | **0.845** | 0.070 | -0.006 | 0.015 | 0.428 | **0.840** | 0.110 | 0.025 | 0.024 |
| Eotaxin | 0.304 | **0.718** | 0.227 | 0.079 | -0.016 | 0.500 | **0.517** | -0.072 | 0.090 | 0.107 | **0.497** | 0.494 | -0.072 | 0.016 | 0.078 |
| IL-8 | 0.125 | **0.692** | 0.398 | -0.039 | -0.052 | 0.192 | **0.695** | 0.237 | -0.034 | -0.090 | 0.143 | **0.762** | 0.242 | 0.009 | -0.057 |
| MIP-1β | 0.019 | 0.070 | **0.960** | 0.010 | 0.045 | 0.069 | -0.003 | **0.860** | -0.122 | 0.014 | -0.008 | 0.120 | **0.750** | -0.058 | 0.049 |
| TNF-α | 0.164 | 0.005 | **0.948** | 0.010 | 0.005 | 0.231 | 0.006 | **0.859** | 0.085 | -0.047 | 0.280 | 0.000 | **0.837** | 0.021 | -0.052 |
| MIP-1α | 0.010 | 0.021 | **0.911** | 0.054 | 0.041 | 0.086 | 0.019 | **0.905** | -0.067 | 0.033 | 0.053 | 0.106 | **0.792** | -0.008 | 0.042 |
| IL-1β | 0.392 | 0.131 | **0.836** | -0.058 | 0.034 | 0.287 | 0.072 | **0.762** | -0.170 | -0.043 | 0.202 | 0.067 | **0.751** | -0.131 | -0.041 |
| IL-6 | 0.099 | 0.305 | **0.762** | -0.049 | 0.489 | 0.291 | **0.747** | 0.450 | -0.011 | 0.046 | 0.308 | **0.793** | 0.352 | 0.017 | 0.069 |
| MCP-1 | 0.097 | 0.617 | **0.527** | -0.032 | -0.014 | 0.064 | **0.638** | 0.620 | -0.026 | -0.069 | 0.084 | **0.752** | 0.449 | 0.048 | -0.037 |
| IP-10 | 0.118 | 0.005 | **0.362** | -0.079 | -0.092 | -0.169 | 0.087 | **0.677** | -0.028 | -0.024 | 0.093 | -0.034 | **0.768** | -0.046 | -0.037 |
| PDGF-AA | 0.114 | 0.021 | 0.018 | **0.942** | -0.032 | 0.262 | -0.116 | -0.051 | **0.881** | -0.059 | 0.202 | -0.099 | -0.020 | **0.896** | -0.018 |
| sCD40L | 0.089 | -0.066 | -0.058 | **0.928** | -0.032 | 0.165 | -0.168 | -0.065 | **0.893** | -0.054 | 0.097 | -0.157 | -0.060 | **0.915** | -0.040 |
| PDGF-AB/BB | 0.065 | -0.003 | -0.045 | **0.928** | -0.032 | 0.142 | -0.080 | -0.081 | **0.913** | -0.051 | 0.031 | -0.048 | -0.059 | **0.920** | -0.049 |
| GRO | 0.071 | 0.030 | -0.007 | **0.902** | -0.026 | 0.160 | 0.192 | -0.039 | **0.826** | -0.070 | 0.074 | 0.263 | -0.021 | **0.851** | -0.029 |
| EGF | 0.156 | 0.430 | -0.057 | **0.731** | 0.022 | 0.140 | 0.237 | -0.112 | **0.700** | 0.103 | 0.173 | 0.259 | -0.145 | **0.718** | 0.075 |
| RANTES | -0.200 | -0.100 | -0.033 | **0.387** | 0.326 | -0.250 | -0.067 | 0.040 | **0.457** | 0.294 | **-0.397** | 0.034 | -0.044 | 0.371 | 0.294 |
| IL-1α | 0.090 | 0.147 | -0.012 | -0.034 | **0.962** | 0.023 | 0.089 | -0.007 | 0.005 | **0.951** | 0.032 | 0.059 | -0.015 | -0.018 | **0.965** |
| IL-9 | 0.213 | 0.219 | -0.004 | -0.047 | **0.931** | 0.099 | 0.119 | -0.021 | -0.020 | **0.951** | 0.109 | 0.110 | -0.040 | -0.021 | **0.950** |
| IL-4 | 0.317 | 0.155 | 0.129 | 0.008 | **0.861** | 0.179 | 0.107 | -0.026 | 0.042 | **0.882** | 0.188 | 0.082 | 0.059 | 0.037 | **0.917** |

### 3.4. Relationships between Adrenergic Immune Modulation and Psychiatric Measures

Across the three experimental conditions, a similar immune factor (major loadings: PDGF-AB/BB, sCD40L, PDGF-AA, GRO, EGF, + RANTES for Control and LPS conditions, + G-CSF for LPS condition; see **Supplementary Table 7**) was identified to be significantly correlated with anticipatory anhedonia after Bonferroni correction 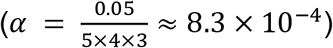 and controlling for age, sex, and BMI. Specifically, correlations were found between anticipatory anhedonia and a) Control Factor (F) 4 (**Figure 1A**, *rho* = −0.35, *p* = 3.6 × 10^−4^), b) LPS F4 (**Figure 1B**, *rho* = −0.36, *p* = 2.6 × 10^−4^), and c) LPS+CBL F4 (**Figure 1C**, *rho* = −0.39, *p* = 7.4 × 10^−5^). In addition, LPS+CBL F4 significantly correlated with consummatory anhedonia (**Figure 1D**, *rho* = −0.36, *p* = 3.3 × 10^−4^).

**Figure 1.**
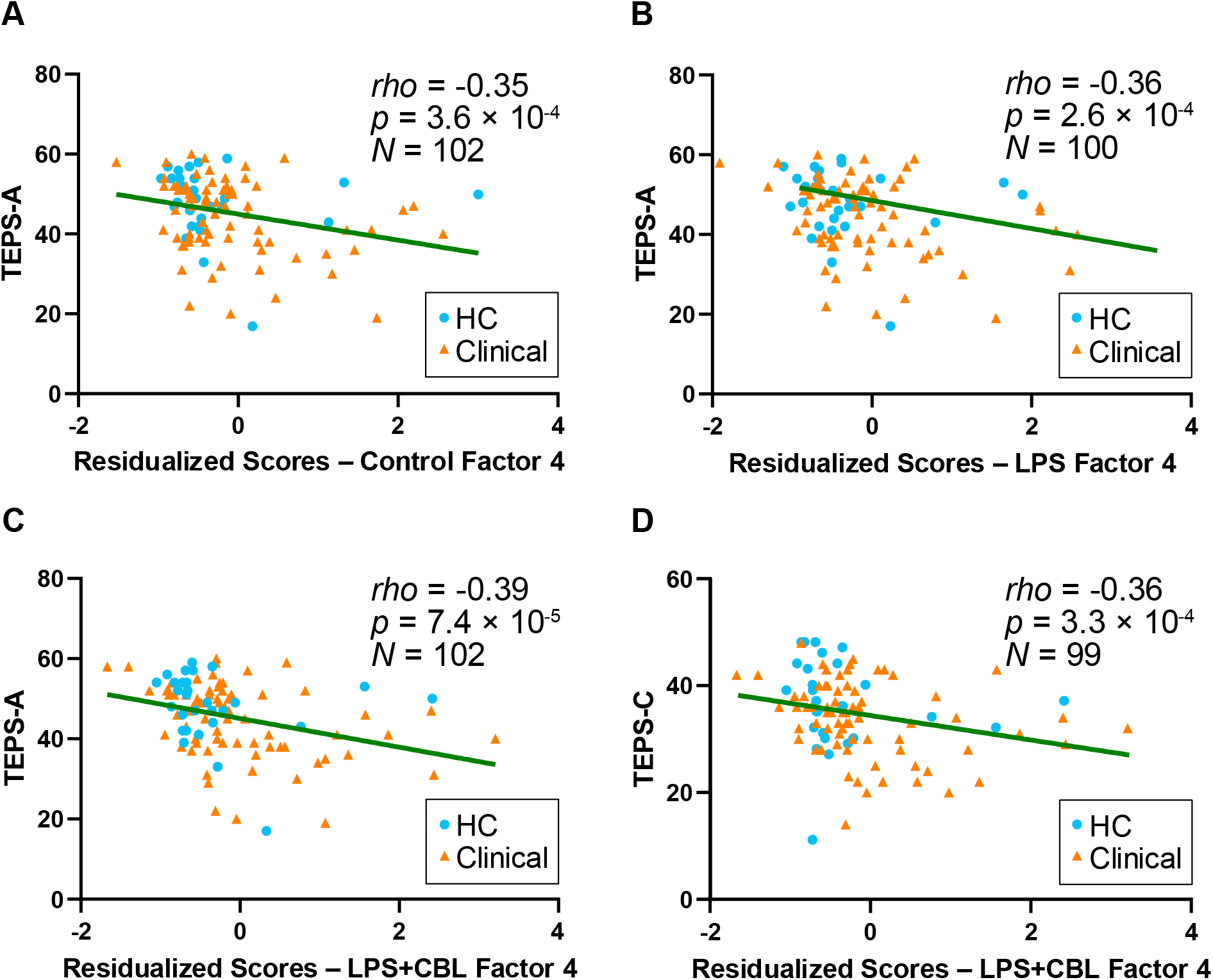
Partial correlations controlling for age, sex, and body mass index (BMI) between **(A)** Control Factor 4 and Anticipatory Anhedonia; **(B)** LPS Factor 4 and Anticipatory Anhedonia; **(C)** LPS+CBL Factor 4 and Anticipatory Anhedonia; **(D)** LPS+CBL Factor 4 and Consummatory Anhedonia. All are significant at Bonferroni-corrected significant threshold 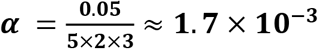. *Abbreviations*: CBL: clenbuterol; HC: healthy control participants; LPS: lipopolysaccharide; Clinical: participants with psychiatric symptoms and/or diagnoses. TEPS-A: Temporal Experience of Pleasure Scale – Anticipatory; TEPS-C: Temporal Experience of Pleasure Scale – Consummatory.

**Figure 2.**
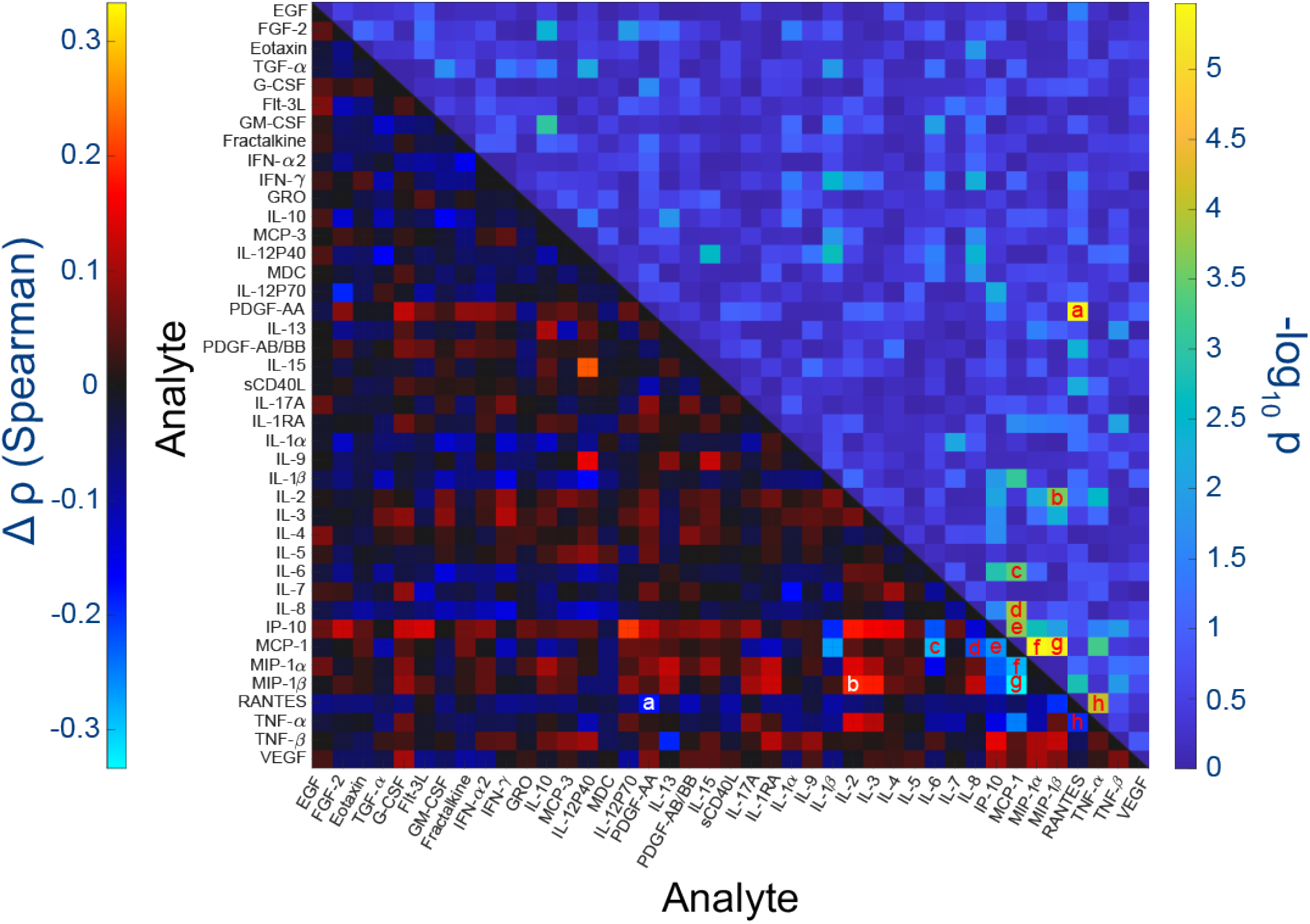
Change in analyte correlations in the LPS+CBL condition relative to the LPS condition (LPS+CBL – LPS; lower half triangle), and their corresponding log-transformed *p* values (upper half triangle). With false discovery rate correction, significant differences were detected in the following pairs: *(a)* RANTES and PDGF-AA, *(b)* MIP-1 β and IL-2, *(c)* MCP-1 and IL-6, *(d)* MCP-1 and IL-8, *(e)* MCP-1 and IP-10, *(f)* MIP-1α and MCP-1, *(g)* MIP-1β and MCP-1, *(h)* RANTES and TNF-α. *Abbreviations*: CBL: clenbuterol; LPS: lipopolysaccharide

When depression severity was also included as a covariate, LPS+CBL F4 remained significantly correlated with anticipatory anhedonia (*rho* = −0.34, *p* = 6.3 × 10^−4^) and consummatory anhedonia (*rho* = −0.34, *p* = 9.9 × 10^−4^), while relationships between anticipatory anhedonia and Control F4 and LPS F4 did not meet the Bonferroni-corrected significant threshold 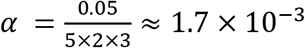 (i.e. adjusting for number of factors, TEPS subscales, and conditions). When these analyses were repeated with total TEPS score (sum of TEPS-A and TEPS-C), these negative correlation trends were similarly observed in immune factors with anhedonia relationships, with LPS+CBL F4 correlating more strongly with anhedonia compared to Control F4 and LPS F4 (see **Supplementary Results –** *SR3*). No significant relationships were found between any immune factor and depression or anxiety. All correlation results are reported in **Supplementary Table 8**.

Across all three conditions, the observed relationships between F4 and anhedonia subconstructs were significantly different from associations between anhedonia and F1, F2, and F3. Additionally, consistent across LPS and LPS+CBL conditions was the significant difference between F4 and F5 in their associations with anticipatory anhedonia. Details of these results can be found in **Supplementary Tables 10 – 11**.

Despite the lack of significant associations between anhedonia scores and analyte ratios between LPS+CBL and LPS conditions (see **Supplementary Results –** *SR1* and **Supplementary Table 9**), our primary results showing differential relationships between anhedonia and factors in these two conditions alone called for further dissection of their immune profiles. The results of our *post hoc* examination of differences in immune profiles across LPS and LPS+CBL conditions were summarized in **Figure 2** and detailed in **Supplementary Results** – *SR4*.

## 4. Discussion

To our knowledge, this study is the first to examine adrenergic modulation of *ex vivo* inflammatory responses using comprehensive immune biomarker assays. As hypothesized, our results indicated that CBL had a general attenuating effect on LPS-induced immune activation in a large sample of medically healthy adolescents with diverse psychiatric profiles. Further, utilizing data-driven factor analysis, we documented consistent relationships between immune protein production and anhedonia in youth across a range of conditions. We discuss our findings below.

### 4.1. Clenbuterol Functionally Modeled Adrenergic Immune Modulation

Using a conservative statistical approach with stringent correction for multiple comparisons, we identified nine immune analytes for which LPS-induced inflammatory responses were significantly attenuated by simultaneous administration of the β_2_-adrenergic receptor agonist CBL, as well as one (sCD40L) for which CBL potentiated release compared to LPS alone. Notably, IL-1β, IL-6, and TNF-α, which have been consistently implicated in psychopathology [2, 51, 52], were among the cytokines most substantially attenuated by CBL in our study. While not previously reported in clinical studies, the observed post-CBL reduced levels of IL-1β, IL-6, and TNF-α mirror results from preclinical studies examining immune activity following CBL induction [53, 54].

Another important finding from our study is the pronounced changes in levels of MIP-1α and MIP-1β chemokines across *ex vivo* conditions. MIP-1α and MIP-1β have been implicated in multiple immune-related pathologies, including those affecting the central nervous system [55]. The detected increase of MIP-1α and MIP-1β levels following LPS stimulation in our study is consistent with previous reports [55]; however, their attenuated levels following CBL addition have never been studied and are novel findings. Though infrequently investigated compared to other immune biomarkers in psychopathologic contexts, MIP chemokines have been assayed in a few studies recruiting adult psychiatric participants [55–57]. One postmortem study found that MIP mRNA was expressed in brain tissue samples of 4 patients with schizophrenia and 1 patient with manic depressive illness but not in those of 8 patients with dementia, neurodegenerative disease, and brain tumor, suggesting a specific role in psychiatric vs. neurological illness [58]. The handful of clinical studies to examine MIP levels in psychiatric cohorts have yielded inconsistent findings, with some reporting elevated serum MIP-1β concentrations in patients with more severe depression [56, 57], but others reporting lower or no difference in MIP levels in patients with major depression compared to healthy controls [59, 60]. Our study presents new evidence supporting MIP chemokines as part of the general immune response to adrenergic stimulation in youth with diverse mood and anxiety symptoms.

While CBL attenuated the release of multiple pro-inflammatory cytokines, it had minimal impact on anti-inflammatory biomarkers, including IL-4 and IL-10 [61, 62]. Animal literature has provided conflicting evidence regarding adrenergic modulation of IL-10, reporting both suppression [25, 63] and induction of IL-10 following administration of β_2_ receptor agonists, including CBL [64, 65]. As IL-4 and IL-10 were elevated in both LPS and LPS+CBL conditions compared to Control in our study, one interpretation is that these cytokines were already being produced in response to LPS to limit the predominantly inflammatory effects of other induced cytokines and thus were not sensitive to the coadministration of CBL. Furthermore, CBL exposure led to increased levels of sCD40L, which is a signaling molecule typically characterized as pro-inflammatory but promotes the secretion of IL-10 and exhibits immunosuppressive activity in HIV infection and certain cancers [66, 67]. In section *4.2* below, we discuss mechanisms possibly underlying CBL’s immune-attenuation effects given the observed minimal change in IL-10 levels between LPS and LPS+CBL conditions in this study. We note that these types of complex interactions between pro- and anti-inflammatory mechanisms are characteristic of the immune system, resulting in varied observations depending on the methods used.

Similarly, several hematopoietic growth factors (i.e. G-CSF, GM-CSF, PDGF-AA/BB, TGF-α) responsive to LPS stimulus did not appear to be strongly affected by CBL. Few studies have examined if and how hematopoietic growth factors expression react to CBL [68, 69], and virtually none addressed such questions in post-endotoxin exposure settings. The lack of substantial change observed peripherally in levels of hematopoietic growth factor following CBL induction could be explained by hematopoietic growth factors acting more centrally in psychopathologic contexts [70].

### 4.2. Mechanisms Underlying Anti-inflammatory Effects by β_2_-adrenergic Signaling

Though β_2_ agonism induces a generalized anti-inflammatory effect, responses to β_2_ signaling by subtypes of immune cells are nuanced [21, 71]. Innate immune cells with well-differentiated functions appear to be consistent in their anti-inflammatory responses to β_2_-adrenergic receptors (AR) activation. Human monocytes primarily exert an immune-suppressive effect through reducing levels of post-LPS MIP-1α, IL-18, and IL-12 production [72, 73]. Dendritic cells, though less commonly studied, have also been shown to primarily engage in anti-inflammatory activities following adrenergic modulation [21].

However, in the absence of inflammatory triggers, β_2_-AR activation might signal monocytes to upregulate pro-inflammatory activities, with increased secretion of IL-18, IL-6, and free radicals [74, 75]. Adaptive immune cells express β_2_-AR differentially, which was thought to influence their responses to post-inflammatory adrenergic stimulation [71, 76]. Naïve CD4+ T cells can differentiate into Th1 or Th2 cells in response to β_2_-AR engagement depending on their cellular environment, and downstream increased IFN-γ production through mature Th1 cells has been reported [77]. B cells, known for their antibody producing capabilities, can initiate a downstream pro-inflammatory effect upon adrenergic stimulation [78].

Beyond the initial responses to adrenergic stimulation, proteins released from these activated cells trigger feedback pathways to regulate the overall production of immune-modulating signaling molecules. Cross-regulations between innate and adaptive branches of the immune system add further granularity to cellular responses following β_2_-agonism [21, 71, 76]. Conflicting reports of various immune protein levels being promoted or suppressed after β_2_-AR stimulation emphasize the complexity of various immune modulating systems and the importance of more global approaches to examine a wide spectrum of immune biomarkers. In our study in which analyte levels were quantified following *in vitro* experimentation on whole blood samples, the reported adrenergic attenuation of immune activation likely reflected the net effect of these mechanisms.

Although we cannot make direct inferences about precise molecular mechanisms of CBL’s downregulating effect on LPS-induced immune dysregulation from the results of this study, intracellular cyclic adenosine monophosphate (cAMP) level elevation might mediate such observed effect, as supported by an *in vitro* study of cellular events in human peripheral blood mononuclear cells and cumulative preclinical studies on mechanisms of β_2_-AR signaling [54, 79]. CBL has also been shown to suppress LPS-induced nuclear factor kappa-light-chain-enhancer of activated B cells (NF-κB) activity through prevention of nuclear factor of kappa light polypeptide gene enhancer in B-cells inhibitor (IκB) phosphorylation in rodent brains [80]. While the mechanism underlying CBL-driven production of hematopoietic growth factors is unclear, it is plausible that NF-κB promotes their secretion and activities [81, 82]. Given that transcription factor NF-κB is ubiquitous in human cells [83], CBL possibly exerts its inflammatory-attenuating property through this modulator, similar to other β_2_-AR agonists [84].

### 4.3. New Insights into Immunologic Correlates of Anhedonia

Our secondary analytic strategy revealed associations between anhedonia sub-components and immune measures at baseline as well as in response to endotoxin and adrenergic challenge. This finding emphasizes the advantages of employing the RDoC approach focusing on behavioral constructs and neuronal circuits rather than limiting investigations to categorical classification of psychiatric disorders. Additionally, this finding fits our prior research reporting relationships between levels of various inflammatory mediators in the blood and anhedonia, as well as their associations with brain function during a reward fMRI task [6, 18–20]. The lack of overlap between immune biomarkers associated with anhedonia in this work and those in our prior work is most likely due to differences in sample size and anhedonia measure (TEPS vs SHAPS). Complementing our works focusing on adolescence, others have shown a correlation between an LPS-induced immune cluster and anticipatory anhedonia, indexed by TEPS-A and a reward fMRI task, in adults [85]. The varied analytes contributing to different immune factors linked to anhedonia across studies is expected from the data-driven nature of factor analyses.

Interestingly, specific to post-CBL exposure, a cluster composed predominantly of hematopoietic growth factors was linked to higher degrees of both anticipatory and consummatory anhedonia. These relationships persisted after adjusting for depression severity. Previously, several growth factors, including G-CSF, GM-CSF, sCD40L, and EGF, have been implicated in depression [86–90]. Our findings corroborated this growing literature and encouraged the utilization of dimensional measures of reward constructs in immunopsychiatric investigations beyond conventional disorder categories. While anticipatory and consummatory anhedonia quantified with clinical self-reports might share some variance, our study supported their divergent immunological correlates across various experimental conditions. Perhaps the distinction between anhedonia sub-components is more apparent biologically, as suggested by works from our group and others [18, 85, 91]. While the effects of β_2_-agonism on hematopoietic growth factors remains mostly undiscerned, CBL might enhance these analytes to act in a compensatory manner, leading to an overall immunosuppressive effect. In the context of our study, it is possible that the signaling cascades involving these proteins were more reactive in participants with higher levels of anhedonia.

Although there was a lack of association between anhedonia and relative changes of immune biomarker levels across LPS and LPS+CBL conditions, in dissecting the inter-analyte relationships within their immune profiles, we showed that the complex interplay between pro-inflammatory, chemokines, and hematopoietic growth factors was potentially responsible for the newly identified factor-anhedonia associations. Taken together, our findings lent support to the developing notion that immunological involvement in psychopathology might not be specific to a group of selected few biomarkers [92, 93].

### 4.4. Clinical Implications of Adrenergic Modulating Interventions in Depression

Our results suggest that targeting β_2_-adrenergic receptors might be a promising anti-inflammatory approach for interventions in a subset of patients suffering from depression, as β_2_-agonism could dampen the effects of peripheral inflammation modulating psychiatric symptoms. This approach should be cautiously considered as attempts to develop adrenergic-specific psychopharmacologic therapy targeting depression have been met with cardiovascular and electrolytic challenges [94], and data to date have yet to suggest a clinically meaningful effect of beta-blockers in treating anxiety disorders [95]. One previous trial of CBL as a potential anti-depressant treatment was conducted in 5 female patients with major depression, and all patients experienced systemic side effects, prompting the authors to conclude CBL was not a viable treatment for depression [96]. Alternatively, previous reports have linked improvement in anhedonia to existing interventions that engage sympathetic and parasympathetic systems, including deep breathing exercises, meditation, and physical exercises [97–100]. This study thus provides early neurobiological evidence to support the effectiveness of such interventions. Relatedly, while TNF-α antagonist infliximab, an anti-inflammatory biologic, failed to produce significant antidepressant effects in patients with major depressive episodes, meaningful improvements in anhedonia were observed [101, 102]. In line with these reports, our findings advocate for anti-inflammatory trials specifically targeting anhedonia.

### 4.5. Limitations and Future Directions

This study has several limitations. First, as analyte levels in our experiment were measured at only one time point (6 hours post-exposure), temporal analyte behaviors following treatment of LPS with and without CBL remain unclear. Additionally, we did not conduct parallel ELISA assays for cross-validation purposes. Future research should measure immune protein levels at several post-exposure time points and incorporate ELISA cross-validation to gain more comprehensive insights into CBL-affected immune analyte behaviors. Second, with our current methods, we cannot make further inferences regarding underlying neural mechanisms that might modulate relationships between adrenergic-induced immune suppression and anhedonia. Since we also collect neuroimaging data on many of these participants as part of our on-going research program, we aim to incorporate measurements of reward dysfunctions in our future studies to examine CBL-affected immune response and reward-related neural activity. Third, our study design is cross-sectional, limiting our ability to infer stability of these associations over time. Our analysis also did not account for several factors influencing the immune system and processes that modulate it, including menstrual cycle stage, exercise, diet, stress, and sleep. These data should be collected in future studies to further control for latent confounders and parse out more specific relationships between immune attenuation and symptomatology in adolescents. Finally, the supernatants collected following cell cultures were stored at −20°C, whereas −80°C would be preferable for long-term storage. However, we showed that storage time did not influence measured levels of 40 out of 41 analytes in our study, supporting the validity of our findings.

### 4.6. Conclusions

The results of this study further inform the research and clinical communities about CBL’s anti-inflammatory effects that can potentially target neuroinflammation implicated in adolescent psychopathology. More broadly, the results of this study can pave the way for further understanding of the role of β_2_-agonism underlying reward-related constructs that contribute to mood and anxiety symptomatology.

## Author Contributions

Conceptualization: T.N, B.E., S.K, and V.G.

Data Curation: T.N., B.E., D.P, M.P, H.X., S.K., and V.G.

Formal Analysis: T.N.

Funding Acquisition: T.N., B.E, and V.G.

Investigation: T.N., B.E., M.P, H.X., S.K., and V.G.

Methodology: T.N., B.E., M.P, H.X, S.K, and V.G.

Project Administration: T.N., B.E., S.K., and V.G.

Resources: S.K and V.G.

Software: T.N. and B.E.

Supervision: V.G.

Validation: T.N, B.E., S.K., and V.G.

Visualization: T.N.

Writing – Original Draft: T.N., B.E., S.K., and V.G.

Writing – Review & Editing: T.N, B.E., D.P, M.P, H.X., S.K., and V.G.

