## Supplementary Materials for "Clenbuterol Attenuates Immune Reaction to Lipopolysaccharide and Its Relationship to Anhedonia in Adolescents"

#### Supplementary Methods

##### *SM1. Sample Processing and Experimental Conditions*

ACD-anticoagulated whole blood (WB) was collected. WB was then mixed 1:1 with the culture medium, aliquot 950 µL per well in 96-deep-well plates in duplicates. For the Control condition, an additional 50 µL of culture medium was added. Prior to being added to the WB + Medium samples, LPS stock of 1 mg/mL was first diluted 1:10 to 100 µg/ml, then further diluted 1:50 to 2 µg/mL; CLB stock of 0.1 M was diluted 1:100 to $10^{-3}$ M. For the LPS condition, 50 µL of 2 µg/mL LPS was added to the prepared 950 µL WB + Medium sample. The final working concentration of LPS in this culture condition was 0.1 µg/mL. For the LPS+CBL condition, 49 µL of 2 µg/mL LPS and 1 µL of $10^{-3}$ M CBL was added to the prepared 950 µL WB + Medium sample. The final working concentrations of LPS and CBL in this culture condition were 0.1 µg/mL and $10^{-6}$ M, respectively.

The plates were subsequently incubated for 6 hours at 37°C in a 5% $CO_2$ tissue culture incubator. Following incubation, the plates were spun down at 2000 g for 15 minutes at 4°C. Then, 200 µL of cultured supernatant was collected from each well, transferred to PCR-capped tube strips, and stored in a -20°C freezer. The remaining supernatant was removed. The 96-deep-well plates left with whole blood pellet were sealed with well covers and also stored in the -20°C freezer.

##### *SM2. Immune Biomarker Analytes*

Samples from 130 participants were assayed across 21 plates. Acquired fluorescence data were analyzed by the Beadview software (Upstate Inc., Charlottesville, VA). In each plate, analyte levels were determined in duplicate from 25 µL volumes of supernatant collected from cultured whole blood samples using the multiplex panel (MILLIPLEX MAP Human Cytokine/Chemokine Magnetic Bead Panel - Premixed 41 Plex, Millipore Corp., Burlington, MA) as per manufacturer’s instructions. The kit, which included lyophilized pre-mixed standard, was reconstituted and used to generate 6-point serial dilutions. The dilutions were run on each plate along with the samples. There was no dilution of samples on Luminex assays. The culture supernatants were run neat on Luminex assays, and no dilution factor was applied to LPS-stimulated samples.

Analyte median fluorescent intensity (MFI) values were measured twice (once per duplicate sample) and averaged to obtain detailed subject-level immune profiles. The MFI values across samples were automatically generated from bead counts and fluorescent intensity based on the standard instructions from the manufacturer. Any bead counts lower than 35 would prompt further quality control checks. Any final bead count of 0 would result in missing MFI value (uncomputable). To account for potential assay drifts, we conducted normalization between runs to correct for inter-plate batch effects. We used the highest limit of detection (LoD) among all plates to determine the lowest inter-plate LoD. Samples with values below the determined inter-plate LoD were excluded.

Following the manufacturer's instructions, we conducted the following protocol to extract analyte concentration values. In Milliplex Analyst software version 5.1 (Millipore Corp., Burlington, MA), the 5-parameter log fit standard curve for each analyte was generated using: (a) the mean MFI values (from duplicates) obtained from Luminex plate runs, and (b) known concentration for each standard point on the 6-point serial dilutions, as provided by the kit manual. Next, absolute concentration values in pg/mL for unknown samples were determined based on the standard curve's known concentration values. To enhance the reproducibility and transparency of our data, we provided the lower and upper detection limits for each analyte concentrations across all plates, as well as the number of samples with detectable values in **Supplementary Table 1** below. Various samples had analyte levels outside of the detectable range, most notably in the Control condition. This is physiologically expected. Additionally, as concentration data were not available for a substantial number of participants, we opted to utilize MFI for our main analyses to ensure proper statistical power. Immunological data of a subset of participants were previously analyzed and reported in Freed *et al* (1, LPS condition only) and Bradley *et al* (2, Control condition only).

Whole blood samples were collected between May 1, 2013, and June 28, 2019. Following incubation in three *in vitro* conditions (Control, LPS, LPS+CBL), the post-centrifuged supernatant was stored at -20°C. Immune marker analyses were conducted in batches between December 1, 2015, and Aug 29, 2019. The median time to analysis was 279 days (Q1: 99 days, and Q3: 499 days). Please see **Supplementary Figure 4** illustrating the distribution of storage time in days.

Due to the differences in storage time across samples, we conducted secondary analyses employing robust regression to determine if storage time affected immune marker levels. Models included MFI level of each analyte as the dependent variable and storage time in days as the predictor. Results indicated that only Flt-3L was significantly affected by storage time (Control: $\beta = 0.012$, $p = 3.63 \times 10^{-5}$; LPS: $\beta = 0.012$, $p = 8.93 \times 10^{-5}$; and LPS+CBL: $\beta = 0.010$, $p = 6.16 \times 10^{-4}$) after Bonferroni correction ($\alpha = \frac{0.05}{41}$). As Flt-3L was not implicated in any of our key findings, such result suggested that our storage protocol did not meaningfully affect our study.

*SM3. Missing Data*

In our study, one participant had missing MFI data for 9 analytes (fractalkine, G-CSF, IFN-α2, IL-7, IL-9, IL-13, MCP-1, MIP-1α, and PDGF-AB/BB) in the control and LPS conditions. A second participant had missing MFI data for 2 analytes (IL-7 and IL-13) in the LPS condition, and a third participant had missing MFI data for 1 analyte (IL-7) in the LPS condition; complete immune profiles were obtained for all other subjects and conditions. For each analysis, participants with incomplete data were excluded if and only if they were missing data required for that particular analysis.

#### Supplementary Results

*SR1. Factor analysis on changes in cytokine levels between LPS and LPS+CBL*

As a follow-up analysis in examining correlations between anhedonia and change in cytokines between LPS and LPS+CBL conditions, we applied the same exploratory factor analysis method as outlined in our **Methods** to the ratio between cytokines of LPS and LPS+CBL conditions. Instead of computing the delta, we opted to use the ratio between LPS+CBL and LPS analyte levels to determine the change due to high inter-subject variability in absolute counts between the 2 conditions. Such approach yielded 4 Ratio factors explaining 65.1% variance of the data. The Ratio factors' major loadings, shown below, were consistent with those of the immune factors from the Control, LPS, and LPS+CBL conditions.

- Ratio Factor (F) 1: IL-2, IL-12P40, TNF-β, IL-13, IL-15, IL-10, IFN-γ, IL-5, IFN-α2, IL-9, IL-12P70, G-CSF, MCP-3, IL-1α, IL-3, GM-CSF, Fractalkine, IL-7
- Ratio F2: MIP-1α, MIP-1β, IL-6, IL-1β, IL-1RA, TNF-α, IP-10, MCP-1, TGF-α, IL-8
- Ratio F3: VEGF, FGF-2, IL-4, IL-17A, Flt-3L, EGF, Eotaxin
- Ratio F4: PDGF-AA, GRO, RANTES, PDGF-AB/BB, sCD40L, MDC

These Ratio factors were then correlated with TEPS-A and TEPS-C scores using Spearman correlations adjusting for covariates, similar to correlation analyses originally conducted for the immune factors of the Control, LPS, and LPS+CBL conditions. All *rho* coefficients and *p*-values can be found in **Supplementary Table 9**. At the uncorrected level of $p < 0.05$, Ratio F3 showed significant association with anticipatory anhedonia adjusting for age, sex, BMI, and depression. After correcting for multiple comparisons with the Bonferroni method ($\alpha = \frac{0.05}{4} = 0.0125$), there was no significant associations.

*SR 2. Comparing strengths of factor-anhedonia associations with bias-corrected accelerated bootstrap*

Following our newly identified associations between immune markers and anhedonia subconstructs, we determined whether the strengths of these relationships were significantly different from each other. Similar to our original method, we calculated pairwise differences (Δ*rho*) between factor-anhedonia associations by z-transforming *rho* coefficients, subtracting them from each other, then transforming them back to *rho* coefficients. Utilizing the same data-driven bias-corrected and accelerated bootstrap with $10^5$ resamples, *p*-values were computed to determine whether the differences in strengths of factor-anhedonia associations (Δ*rho*) were significant. Results were corrected for multiple comparisons at the symptom association level using the Bonferroni method (i.e. 10 pairs per symptom; significant at α = $\frac{0.05}{10} = 0.005$). With the same bootstrapping technique, we generated and compared 95% confidence intervals for *rho* coefficients of factor-symptom associations.

As expected, results from the two analytical approaches were comparable, providing complementary validation. Overall, these results showed a similar pattern across all conditions; the observed relationships between F4 and anhedonia subconstructs were significantly different from associations between anhedonia sub-constructs and F1, F2, and F3. Additionally, consistent across LPS and LPS+CBL conditions was the significant difference between F4 and F5 in their associations with both subcomponents of anhedonia. Full results can be found in **Supplementary Tables 10 – 11**.

*SR 3. Correlating cytokine factors with total TEPS scores.*

As TEPS-A and TEPS-C scores are quite correlated in our sample, we additionally performed *post-hoc* Spearman partial correlations to examine associations between factors and total anhedonia scores (TEPS-T; combining TEPS-A and TEPS-C) adjusting for age, sex, and BMI.

Total anhedonia was found to be more strongly correlated with LPS+CBL F4 (*rho* = -0.40, $p = 6.60 \times 10^{-5}$) compared to LPS F4 (*rho* = -0.36, $p = 4.47 \times 10^{-4}$) and Control F4 (*rho* = -0.34, $p = 6.16 \times 10^{-4}$). When additionally adjusting for depression severity (BDI), the same trend was observed, with stronger correlation between total TEPS scores and LPS+CBL F4 (*rho* = -0.35, $p = 5.24 \times 10^{-4}$)

compared to Control F4 (*rho* = -0.30, $p = 3.00 \times 10^{-3}$) and LPS F4 (*rho* = -0.30, $p = 3.50 \times 10^{-3}$). Such results are consistent with those reported in our original analyses considering TEPS-A and TEPS-C separately, with LPS+CBL F4 showing stronger correlation with anhedonia compared to Control F4 and LPS F4.

*SR 4. Exploring differences in immune profiles between LPS and LPS+CBL conditions*

To compare immune profiles between LPS+CBL and LPS conditions, we conducted a *post hoc* analysis examining potential cross-condition differences in inter-analyte correlations. Details of our analytic plan was described in **Methods** – *2.5.* and the corresponding findings were summarized in **Results** – *3.4*.

Using stringent Bonferroni correction ($\alpha = \frac{0.05}{820} \approx 6.1 \times 10^{-5}$), we found that administration of CBL significantly affected 3 inter-analyte correlations: RANTES and PDGF-AA (Δ*rho* = -0.18, $p = 3.36 \times 10^{-6}$); MIP-1α and MCP-1 (Δ*rho* = -0.29, $p = 3.5 \times 10^{-6}$); and MIP-1β and MCP-1 (Δ*rho* = -0.33, $p = 4.74 \times 10^{-6}$). Using false discovery rate (FDR) correction [51], we identified CBL-induced changes in correlations between 5 additional analyte pairs: MIP-1β and IL-2 (Δ*rho* = 0.19, $p_{FDR} = 0.030$); MCP-1 and IL-6 (Δ*rho* = 0.29, $p_{FDR} = 0.036$); MCP-1 and IL-8 (Δ*rho* = -0.22, $p_{FDR} = 0.021$); MCP-1 and IP-10 (Δ*rho* = -0.26, $p_{FDR} = 0.028$); and RANTES and TNF-α (Δ*rho* = -0.20, $p_{FDR} = 0.018$).

#### Supplementary Figures and Tables

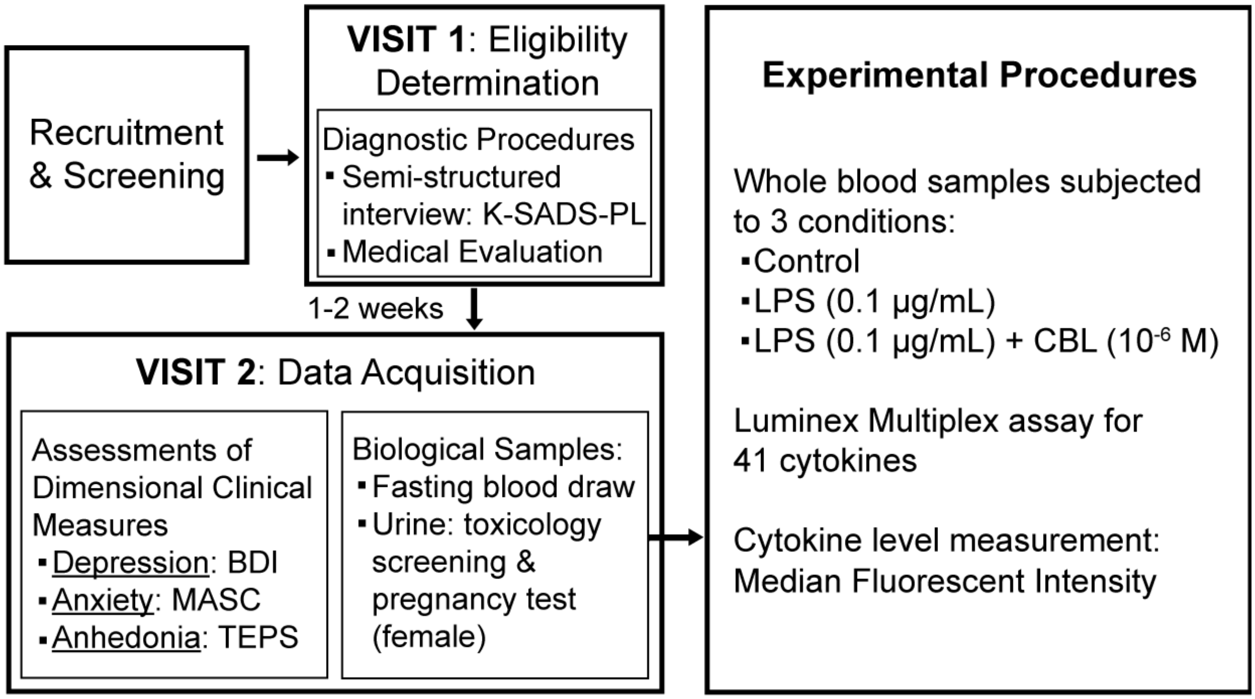

**Supplementary Figure 1**. Summary of study procedures.
*Abbreviations*: BDI: Beck Depression Inventory; CBL: Clenbuterol; K-SADS-PL: Schedule for Affective Disorders and Schizophrenia – Present and Lifetime version; LPS: Lipopolysaccharide; MASC: Multidimensional Anxiety Scale for Children; TEPS: Temporal Experience of Pleasure Scale.

| Full Name | Abbreviation | Class of Immune Marker | Concentration (pg/ml) | | | | | Number of samples with detectable levels | | |
|---|---|---|---|---|---|---|---|---|---|---|
| | | | Lower Detection Limit | Upper Detection Limit | Median (IQR) | | | | | |
| | | | | | Control | LPS | LPS+CBL | Control | LPS | LPS+CBL |
| Epidermal growth factor | EGF | Growth factor | 3.12 | 5608 | 22.76 (14.3 – 41.95) | 23.86 (13.09 – 45.18) | 24.67 (13.65 – 52.37) | 106 | 108 | 109 |
| Eotaxin | Eotaxin | Chemokine | 28.09 | 12519 | 30.74 (17.21 – 48.21) | 31.13 (16.14 – 50.81) | 31.13 (16.09 – 45.31) | 128 | 130 | 130 |
| Fibroblast growth factor-2 | FGF-2 | Hematopoietic growth factor | 19.94 | 9938 | 36.81 (24.88 – 54.9) | 40.64 (24.41 – 56.84) | 39.38 (24.58 – 56.18) | 126 | 126 | 126 |
| FMS-like tyrosine kinase 3-ligand | Flt-3L | Hematopoietic growth factor | 3.83 | 16130 | 14.75 (10.57 – 18.67) | 15 (10.5 – 18.3) | 14.43 (10.84 – 18.29) | 123 | 124 | 123 |
| Fractalkine | Fractalkine | Chemokine | 23.26 | 9981 | 46.83 (30.59 – 68.6) | 51.55 (36.89 – 73.18) | 51.3 (36.59 – 70.73) | 126 | 126 | 127 |
| Granulocyte-macrophage colony-stimulating factor | G-CSF | Hematopoietic growth factor | 3.43 | 10775 | 10.98 (5.54 – 21.31) | 13.55 (6.3 – 22.18) | 13.83 (6.8 – 23.78) | 119 | 121 | 119 |
| Granulocyte colony-stimulating factor | GM-CSF | Hematopoietic growth factor | 3.06 | 11492 | 3.52 (2.96 – 5.48) | 4.45 (2.9 – 5.73) | 2.96 (2.73 – 5.6) | 29 | 39 | 32 |
| Growth regulated oncogene | GRO | Chemokine | 3.44 | 11383 | 142.34 (58.97 – 416.06) | 151.24 (74.42 – 461.86) | 195.06 (70.35 – 486.41) | 126 | 127 | 127 |
| Interferon-alpha 2 | IFN-α2 | Cytokine | 3.69 | 11766 | 9.48 (5.97 – 13.32) | 10.1 (6.33 – 16.27) | 9.66 (6.16 – 14.3) | 110 | 110 | 114 |
| Interferon-gamma | IFN-γ | Cytokine | 3.56 | 12950 | 7.03 (4.22 – 11.12) | 6.48 (4.16 – 13.24) | 6.75 (4.24 – 11.68) | 93 | 102 | 101 |
| Interleukin-10 | IL-10 | Cytokine | 3.31 | 13039 | 5.78 (3.22 – 10.68) | 6.12 (3.49 – 11.53) | 5.93 (3.47 – 10.26) | 57 | 77 | 76 |
| Interleukin-12P40 | IL-12P40 | Cytokine | 3.41 | 1727 | 5.35 (3.11 – 13.3) | 5.6 (3.58 – 15.67) | 5.77 (3.3 – 12.68) | 89 | 102 | 101 |
| Interleukin-12P70 | IL-12P70 | Cytokine | 3.02 | 12780 | 3.84 (3.09 – 7.44) | 4.05 (3.09 – 6.11) | 3.84 (3.02 – 5.69) | 48 | 57 | 53 |
| Interleukin-13 | IL-13 | Cytokine | 3.04 | 9827 | 11.61 (2.56 – 53.75) | 5.94 (2.96 – 43.02) | 7.31 (3.05 – 39.27) | 55 | 58 | 55 |
| Interleukin-15 | IL-15 | Cytokine | 2.86 | 10888 | 3.68 (2.32 – 6.76) | 3.78 (2.74 – 6.2) | 4.04 (2.27 – 5.76) | 54 | 63 | 61 |
| Interleukin-17A | IL-17A | Cytokine | 2.99 | 11514 | 6.34 (3.82 – 13.32) | 4.23 (3.39 – 10.37) | 3.86 (3.2 – 9.63) | 20 | 23 | 22 |
| Interleukin-1 receptor antagonist | IL-1RA | Cytokine | 3.27 | 12582 | 7.03 (4.23 – 8.93) | 5.15 (3.42 – 7.55) | 5.75 (3.58 – 9.56) | 14 | 16 | 11 |
| Interleukin-1 alpha | IL-1α | Cytokine | 3.08 | 10556 | 4.82 (3.27 – 8.6) | 10.75 (4.2 – 32.89) | 8.06 (4.24 – 21.34) | 27 | 89 | 83 |
| Interleukin-1 beta | IL-1β | Cytokine | 3.06 | 9851 | 11.44 (7.17 – 26.21) | 23.71 (13.27 – 81.43) | 18.5 (11.1 – 50.79) | 119 | 128 | 127 |
| Interleukin-2 | IL-2 | Cytokine | 2.94 | 9544 | 4.71 (3.45 – 7.31) | 4.23 (3.19 – 6.53) | 3.95 (2.89 – 6.63) | 15 | 15 | 13 |
| Interleukin-3 | IL-3 | Cytokine | 3.13 | 2117 | N/A | N/A | N/A | 0 | 0 | 0 |
| Interleukin-4 | IL-4 | Cytokine | 4.67 | 11235 | 11.88 (5.89 – 19.76) | 12.93 (6.77 – 21.25) | 12.44 (6.78 – 24.93) | 91 | 97 | 95 |
| Interleukin-5 | IL-5 | Cytokine | 3.05 | 6265 | 8.77 (4.48 – 25.62) | 9.73 (4.6 – 27.92) | 8.88 (3.71 – 24.53) | 28 | 26 | 28 |

| | | | | | | | | | | |
|---|---|---|---|---|---|---|---|---|---|---|
| Interleukin-6 | IL-6 | Cytokine | 4.62 | 5553 | 10.29 (5.27 – 17.77) | 20.09 (8.85 – 62.48) | 14.36 (7.17 – 35.31) | 35 | 110 | 102 |
| Interleukin-7 | IL-7 | Cytokine | 4.87 | 7535 | 4.45 (2.4 – 7.68) | 4.88 (2.29 – 7.25) | 4.31 (2.81 – 7.06) | 42 | 50 | 42 |
| Interleukin-8 | IL-8 | Chemokine | 3.12 | 6740 | 9.99 (5.08 – 26.41) | 21.68 (9.25 – 47.16) | 15.96 (9.13 – 45.61) | 64 | 121 | 122 |
| Interleukin-9 | IL-9 | Cytokine | 3.04 | 5559 | 4.18 (2.96 – 5.72) | 4.46 (3.01 – 7.19) | 3.88 (3.21 – 6.04) | 16 | 16 | 13 |
| Interferon gamma-induced protein-10 | IP-10 | Chemokine | 21.6 | 13541 | 110.8 (78.92 – 157.66) | 141.89 (95.94 – 237.65) | 119.81 (89.11 – 165.35) | 130 | 130 | 130 |
| Monocyte chemotactic protein-1 | MCP-1 | Chemokine | 4.62 | 5298 | 65.82 (40.87 – 89.85) | 91.05 (56.34 – 139.51) | 76.16 (47.64 – 104.93) | 130 | 130 | 130 |
| Monocyte chemotactic protein-3 | MCP-3 | Chemokine | 4.25 | 12294 | 17.58 (9.18 – 51.57) | 23.17 (10.58 – 57.27) | 18.49 (10.09 – 48.34) | 124 | 124 | 124 |
| Macrophage-derived chemokine | MDC | Chemokine | 16.0 | 10905 | 181.67 (130.66 – 256.68) | 193.3 (129.73 – 274.67) | 188.1 (131.31 – 275.94) | 130 | 130 | 130 |
| Macrophage inflammatory protein-1 alpha | MIP-1α | Chemokine | 9.4 | 821.77 | 12.96 (5.55 – 23.63) | 23.09 (9.16 – 55.92) | 19.4 (8.3 – 31.17) | 46 | 97 | 76 |
| Macrophage inflammatory protein-1 beta | MIP-1β | Chemokine | 3.17 | 1334 | 7.48 (4.55 – 30.62) | 73.04 (32.13 – 169.65) | 34.13 (16.1 – 97.84) | 112 | 130 | 130 |
| Platelet-derived growth factor-AA | PDGF-AA | Hematopoietic growth factor | 16.0 | 990.68 | 86.91 (38.26 – 207.39) | 99.28 (38.98 – 205.62) | 95.76 (45.1 – 225.72) | 129 | 129 | 129 |
| Platelet-derived growth factor-AB/BB | PDGF-AB/BB | Hematopoietic growth factor | 23.64 | 9993 | 929.02 (360.91 – 2670.75) | 1005 (382.37 – 2929) | 1182 (456.22 – 3082.25) | 126 | 123 | 124 |
| Regulated upon Activation, Normal T cell Expressed, and Secreted (or Chemokine ligand 5/CCL5) | RANTES | Chemokine | 51.47 | 11162 | 3428 (1242 – 6478) | 3429 (1295 – 6590) | 4369 (1676 – 6922.5) | 129 | 127 | 127 |
| Soluble cluster of differentiation 40 ligand | sCD40L | Hematopoietic growth factor | 7.24 | 9520 | 84.57 (21.29 – 233) | 78.65 (22.82 – 249.1) | 102.53 (28.68 – 314) | 130 | 129 | 129 |
| Transforming growth factor-alpha | TGF-α | Hematopoietic growth factor | 3.47 | 4048 | 3.92 (2.96 – 6.97) | 3.36 (2.36 – 5.5) | 3.95 (3.02 – 6.2) | 13 | 23 | 15 |
| Tumor necrosis factor-alpha | TNF-α | Cytokine | 3.01 | 9977 | 4.37 (3.22 – 6.54) | 19.81 (9.58 – 39.32) | 10.39 (5.87 – 18.5) | 78 | 125 | 118 |
| Tumor necrosis factor-beta | TNF-β | Cytokine | 3.31 | 9717 | 8.23 (3.78 – 67.14) | 7.75 (3.53 – 63.98) | 8.84 (3.52 – 55.1) | 53 | 50 | 50 |
| Vascular endothelial growth factor | VEGF | Hematopoietic growth factor | 17.16 | 9932 | 37.17 (16.18 – 85.42) | 39.49 (19.07 – 89.23) | 40.04 (19.16 – 91.57) | 115 | 120 | 118 |

**Supplementary Table 1**. List of 41 immune marker analytes, abbreviations, class of immune marker, detection limits, mean, standard deviation, and number of samples with detectable levels in pg/ml.

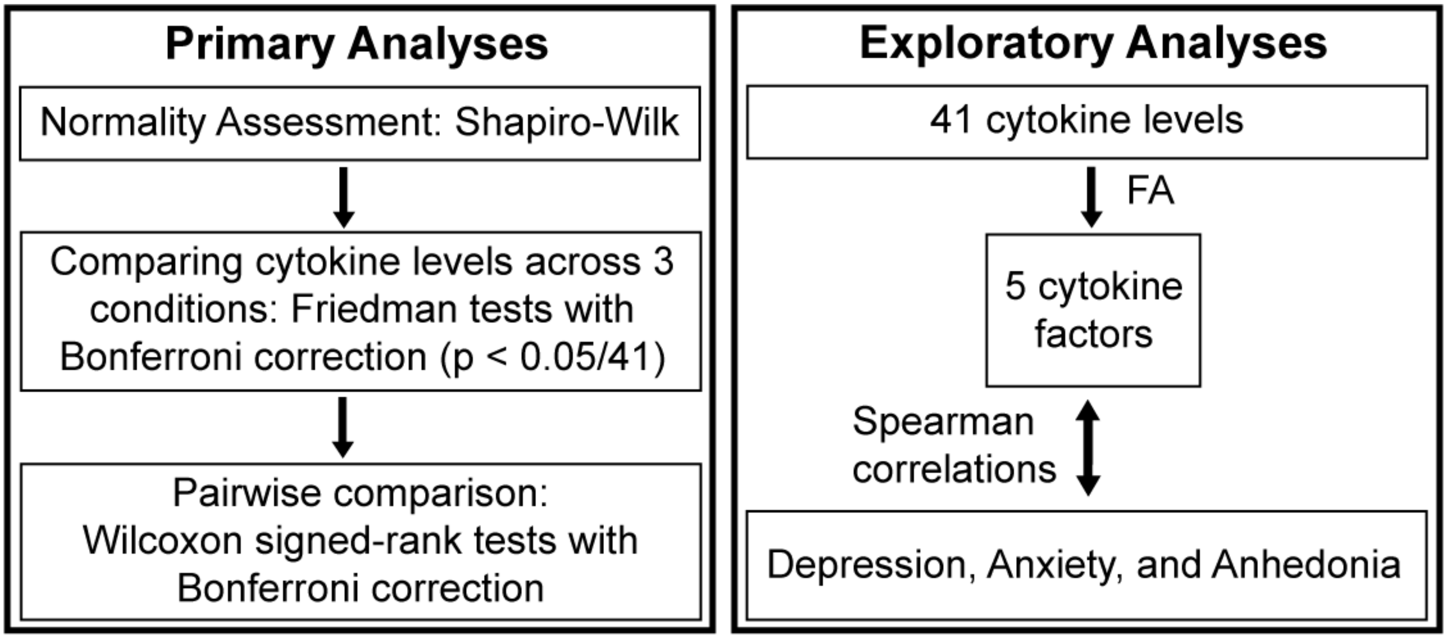

**Supplementary Figure 2**. Summary of statistical analyses conducted in our study.
*Abbreviations*: FA: Factor Analysis.

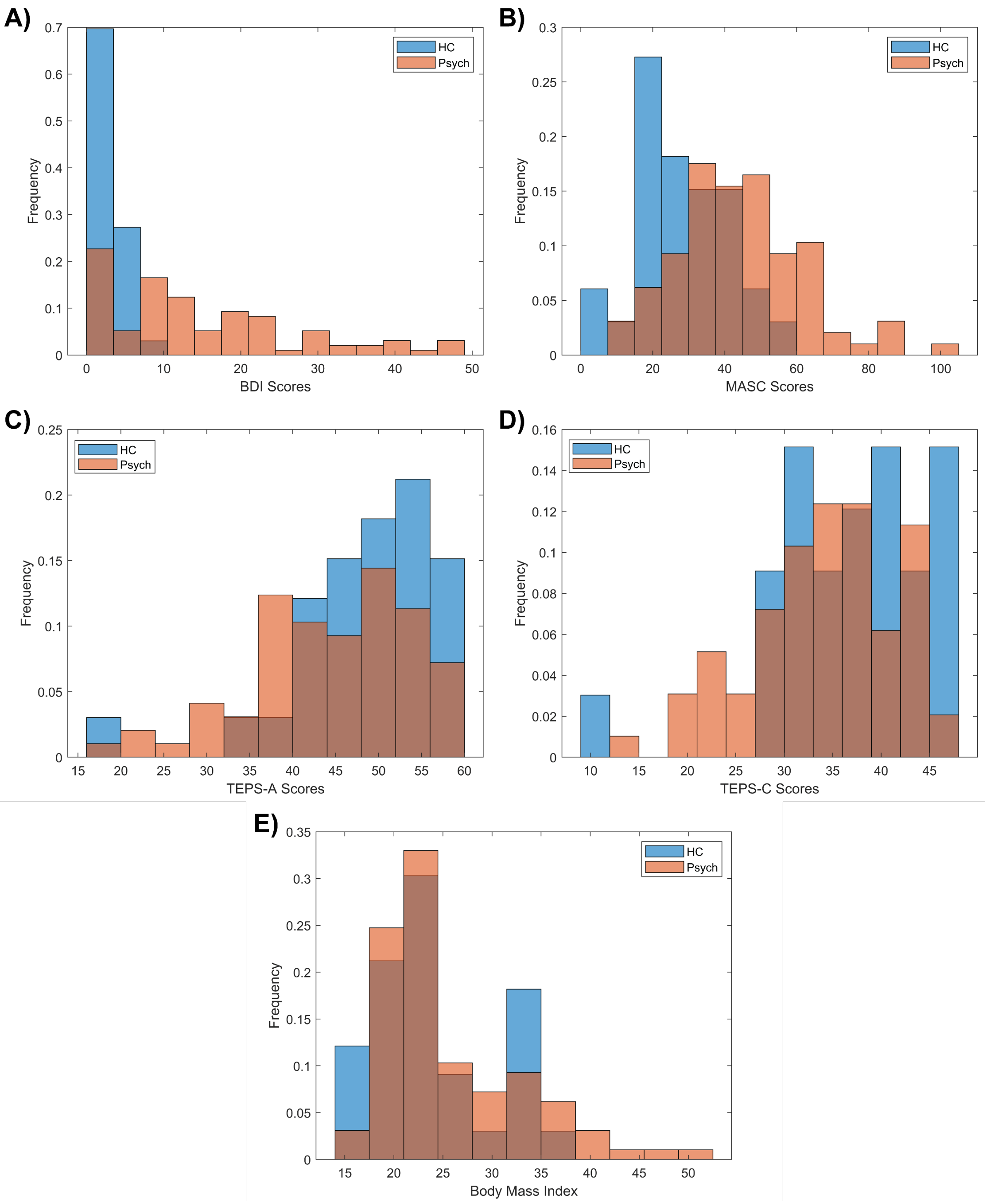

**Supplementary Figure 3**. Distributions of **(A)** depression, **(B)** anxiety, **(C)** anticipatory anhedonia, **(D)** consummatory anhedonia, and **(E)** body mass index (BMI – kg/m$^2$) in our adolescent sample.
*Abbreviations*: BDI: Beck Depression Inventory; HC: healthy control participants; MASC: Multidimensional Anxiety Scale for Children; Psych: participants with psychiatric symptoms and/or diagnoses; TEPS-A: Temporal Experience of Pleasure Scale – Anticipatory; TEPS-C: Temporal Experience of Pleasure Scale – Consummatory.

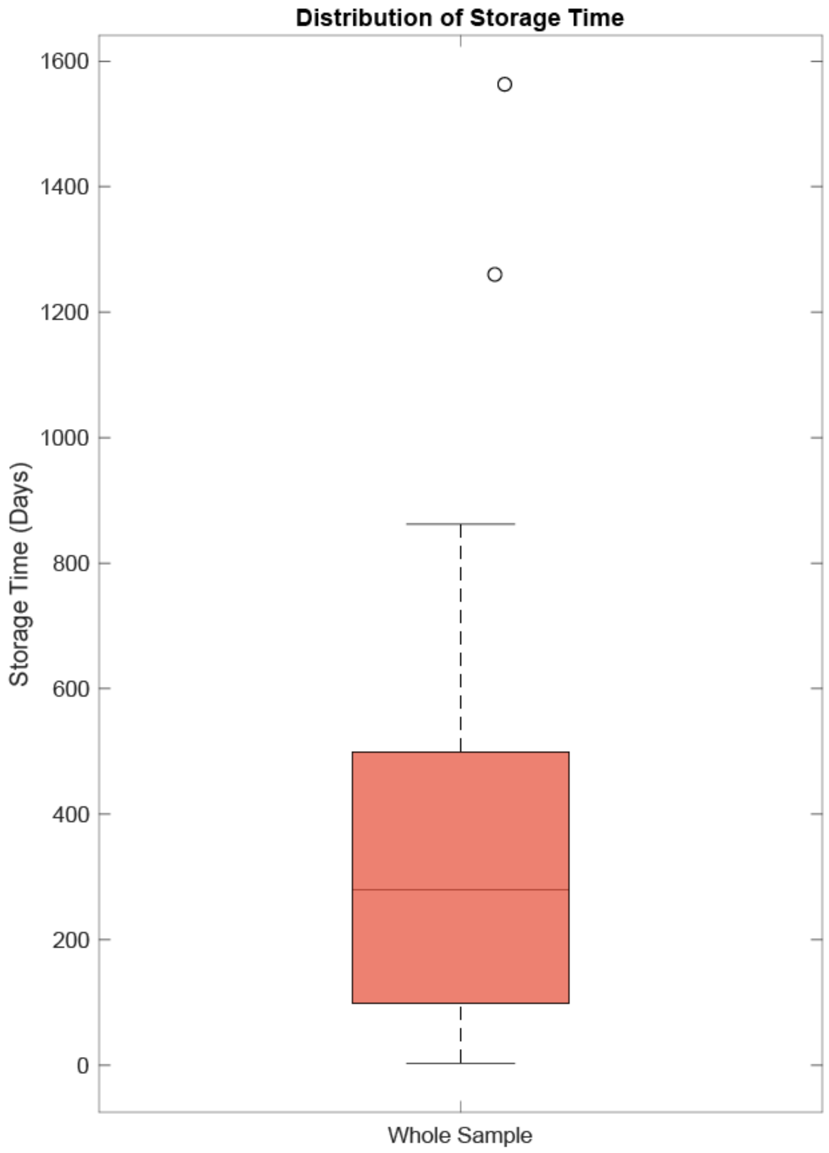

**Supplementary Figure 4**. Box plot showing distribution of sample storage time in days in our whole sample (N=130). Red solid line represents the median. The dots represent outliers.

| | BDI | MASC | TEPS-A | TEPS-C |
|---|---|---|---|---|
| **BDI** | 1 | | | |
| **MASC** | 0.58 ** | 1 | | |
| **TEPS-A** | -0.43 ** | -0.16 | 1 | |
| **TEPS-C** | -0.30 * | -0.12 | 0.49** | 1 |

**Supplementary Table 2.** Spearman correlation matrix of dimensional measures of depression (BDI), anxiety (MASC), anticipatory anhedonia (TEPS-A), and consummatory anhedonia (TEPS-C) in the study sample.
*: $p < 0.05$
**: $p < 0.001$

| Cytokine | Condition | | |
|---|---|---|---|
| | Control | LPS | LPS + CBL |
| EGF | 0.14 | 0.08 | 0.12 |
| Eotaxin | 0.81 | 0.48 | 0.43 |
| FGF-2 | 0.58 | 0.40 | 0.33 |
| Flt-3L | 0.64 | 0.72 | 0.77 |
| Fractalkine | 0.14 | 0.05 | 0.12 |
| G-CSF | 0.24 | 0.28 | 0.23 |
| GM-CSF | 0.13 | 0.04 | 0.07 |
| GRO | 0.09 | 0.06 | 0.04 |
| IFN-α2 | 0.18 | 0.28 | 0.16 |
| IFN-γ | 0.90 | 0.83 | 0.86 |
| IL-10 | 0.03 | 0.02 | 0.02 |
| IL-12P40 | 0.60 | 0.90 | 0.97 |
| IL-12P70 | 0.12 | 0.10 | 0.04 |
| IL-13 | 0.86 | 0.83 | 0.57 |
| IL-15 | 0.59 | 0.74 | 0.47 |
| IL-17A | 0.26 | 0.25 | 0.12 |
| IL-1RA | 0.29 | 0.19 | 0.26 |
| IL-1α | 0.42 | 0.33 | 0.68 |
| IL-1β | 0.11 | 0.04 | 0.07 |
| IL-2 | 0.17 | 0.30 | 0.16 |
| IL-3 | 0.07 | 0.07 | 0.11 |
| IL-4 | 0.89 | 0.90 | 0.74 |
| IL-5 | 0.21 | 0.16 | 0.23 |
| IL-6 | 0.86 | 0.99 | 0.89 |
| IL-7 | 0.15 | 0.24 | 0.17 |
| IL-8 | 0.75 | 0.31 | 0.67 |
| IL-9 | 0.27 | 0.17 | 0.13 |
| IP-10 | 0.34 | 0.30 | 0.52 |
| MCP-1 | 0.22 | 0.34 | 0.26 |
| MCP-3 | 0.93 | 0.91 | 0.94 |
| MDC | 0.58 | 0.86 | 0.97 |
| MIP-1α | 0.17 | 0.34 | 0.30 |
| MIP-1β | 0.55 | 0.68 | 0.97 |
| PDGF-AA | 0.97 | 0.50 | 0.67 |
| PDGF-AB/BB | 0.79 | 0.99 | 0.90 |
| RANTES | 0.34 | 0.83 | 0.64 |
| sCD40L | 0.69 | 0.66 | 0.89 |
| TGF-α | 0.10 | 0.10 | 0.05 |
| TNF-α | 0.41 | 0.45 | 0.68 |
| TNF-β | 0.32 | 0.55 | 0.44 |
| VEGF | 0.52 | 0.31 | 0.28 |

**Supplementary Table 3.** Results of Wilcoxon rank sum tests (*p* values) comparing cytokine levels between participants with psychiatric symptoms and healthy controls. None were significant at $\alpha = \frac{0.05}{41}$ (Bonferroni correction), suggesting no differences in cytokine levels between the two groups across 3 conditions.

| **Cytokine** | **Mean fluorescence intensity** | | | ***p* value** |
|---|---|---|---|---|
| | **Control** | **LPS** | **LPS + CBL** | |
| EGF | 131.5 (58 - 434.5) | 158.125 (56 - 477) | 169 (62.5 - 531.75) | 0.04 |
| Eotaxin | 87 (41.5 - 138.5) | 93.5 (51 - 144.75) | 89.875 (42.75 - 133) | 0.42 |
| FGF-2 | 16.125 (13.5 - 21) | 16.875 (14 - 22) | 16.875 (14 - 22.5) | 0.05 |
| Flt-3L | 25.5 (22 - 32.25) | 26.625 (21 - 33.5) | 26.5 (21.5 - 32.5) | 0.2 |
| Fractalkine | 18.5 (15.5 - 23.8125) | 19.5 (16.25 - 25.0625) | 19.25 (16.1875 - 23.5) | $\mathbf{3.69 \times 10^{-7}}$ ***** |
| G-CSF | 24.5 (18.69 - 35.31) | 27.75 (19.5 - 42.94) | 27 (20.6875 - 36.88) | $\mathbf{8.27 \times 10^{-4}}$ ***** |
| GM-CSF | 21.625 (18 - 27.5) | 23.375 (19.25 - 31.5) | 23.5 (19.25 - 30.25) | $\mathbf{7.95 \times 10^{-6}}$ ***** |
| GRO | 726.25 (171.5 - 3667.8) | 927.125 (253.5 - 4251.5) | 1375.4 (279 - 5024) | $\mathbf{2.26 \times 10^{-8}}$ ***** |
| IFN-α2 | 19.5 (16.69 - 26) | 22 (16.75 - 28.8125) | 21.5 (17 - 29.5) | 0.02 |
| IFN-γ | 24.625 (20 - 34.25) | 26.25 (21.5 - 37) | 25.875 (21 - 37.5) | 0.01 |
| IL-10 | 23.25 (19 - 30.5) | 25.5 (21.5 - 41) | 26.5 (22 - 39.5) | $\mathbf{1.29 \times 10^{-12}}$ ***** |
| IL-12P40 | 24.75 (20 - 34.25) | 27.5 (21 - 40.25) | 26.625 (20.5 - 35.25) | $\mathbf{2.31 \times 10^{-4}}$ ***** |
| IL-12P70 | 17.5 (14.5 - 22) | 18 (15 - 24) | 18.375 (15.5 - 22.5) | $7.72 \times 10^{-3}$ |
| IL-13 | 21 (15.375 - 40) | 22 (16.5 - 39.5) | 23 (16.25 - 39.375) | 0.1 |
| IL-15 | 25.75 (19.75 - 37.5) | 26.875 (20.75 - 38.5) | 27 (21.5 - 40) | 0.06 |
| IL-17A | 28.875 (22 - 38) | 31 (23.25 - 37.5) | 29.625 (23.5 - 38.25) | 0.35 |
| IL-1RA | 24 (18 - 51) | 47.75 (23 - 110.75) | 37.25 (21 - 86.25) | $\mathbf{1.82 \times 10^{-27}}$ ***** |
| IL-1α | 32.5 (24.5 - 40) | 36 (27.5 - 51.25) | 34.5 (28.75 - 43.5) | $\mathbf{1.58 \times 10^{-6}}$ ***** |
| IL-1β | 22.125 (18 - 30.5) | 68.125 (34.5 - 194) | 55.25 (35 - 143.5) | $\mathbf{3.80 \times 10^{-34}}$ ***** |
| IL-2 | 23.125 (18.5 - 54.5) | 26 (18 - 55.5) | 25.5 (18.5 - 56) | 0.17 |
| IL-3 | 17 (12.5 - 24) | 17.5 (13 - 24) | 17.5 (13.5 - 26) | 0.03 |
| IL-4 | 19 (15.5 - 29.25) | 20.875 (17 - 31) | 21.875 (17 - 32.25) | $\mathbf{3.64 \times 10^{-6}}$ ***** |
| IL-5 | 18.5 (16 - 27.75) | 19.5 (16 - 26.5) | 20.25 (16.5 - 28) | 0.11 |
| IL-6 | 28.5 (20.75 - 53) | 165.25 (68.75 - 467.5) | 118.25 (56.5 - 290.5) | $\mathbf{6.42 \times 10^{-38}}$ ***** |
| IL-7 | 16.75 (14.5 – 25) | 19 (15.25 - 27.5) | 19 (15 - 26.5) | $\mathbf{1.82 \times 10^{-6}}$ ***** |
| IL-8 | 86.5 (58 - 226.5) | 466.5 (235.5 - 1195.2) | 435.25 (220.5 - 1201) | $\mathbf{3.92 \times 10^{-33}}$ ***** |
| IL-9 | 29.5 (22.1875 - 43.9375) | 31 (22.1875 - 51.75) | 31.75 (23 - 53) | $4.15 \times 10^{-3}$ |
| IP-10 | 245.5 (168.5 - 360.5) | 337.875 (224.25 - 733) | 271.5 (178 - 430.25) | $\mathbf{3.65 \times 10^{-18}}$ ***** |
| MCP-1 | 652.5 (346.88 - 989.58) | 1030.8 (572 - 1867.78) | 825.25 (448.56 - 1434.05) | $\mathbf{3.45 \times 10^{-19}}$ ***** |
| MCP-3 | 21.5 (15.5 - 62.25) | 25.5 (16.5 - 87) | 24.75 (16 - 78.5) | 0.13 |
| MDC | 438.25 (278.25 - 565.5) | 413.25 (301 - 616.25) | 422.375 (261.5 - 589.75) | 0.43 |
| MIP-1α | 27 (18.5 - 97.25) | 228 (86.75 - 866.3125) | 89.25 (38.4375 - 334.875) | $\mathbf{4.43 \times 10^{-32}}$ ***** |
| MIP-1β | 33.875 (21 - 67) | 356.25 (143.5 - 1102) | 126.875 (58.75 - 465.75) | $\mathbf{3.14 \times 10^{-41}}$ ***** |
| PDGF-AA | 1485.65 (817.75 - 3851.8) | 1684.75 (822.75 - 3759.2) | 1816.9 (873.25 - 4192.20) | 0.14 |
| PDGF-AB/BB | 333 (153 - 1073.1) | 356 (151.625 - 1312.775) | 438.25 (180.06 - 1444.23) | $\mathbf{7.57 \times 10^{-4}}$ ***** |
| RANTES | 9425.75 (3137.8 - 15368) | 9432.25 (3858.8 - 15428) | 10867 (4581.20 - 16561) | $2.58 \times 10^{-3}$ |
| sCD40L | 189.125 (78 - 765.25) | 210.75 (83.25 - 829.5) | 253.5 (104.5 - 1012.5) | $\mathbf{3.36 \times 10^{-6}}$ ***** |
| TGF-α | 23.5 (18.5 - 34) | 28 (22.5 - 49) | 28.25 (21.5 - 42.5) | $\mathbf{8.13 \times 10^{-16}}$ ***** |
| TNF-α | 49.5 (34.5 - 67) | 173 (92 - 456) | 101.625 (56 - 200) | $\mathbf{1.16 \times 10^{-43}}$ ***** |
| TNF-β | 23 (17.5 - 50.25) | 23.5 (18 - 45) | 23.125 (18.5 - 49.5) | 0.65 |
| VEGF | 23 (19.5 - 32.5) | 24.125 (20 - 35.75) | 25.5 (20.5 - 33.25) | $2.42 \times 10^{-3}$ |

**Supplementary Table 4.** Cytokine, chemokine, and growth factor mean fluorescence intensity levels (median and interquartile range) across 3 *ex vivo* conditions in whole sample (N=130). Friedman test *p* values are presented.

*: significant at α = $\mathbf{\frac{0.05}{41} \approx 1.2 \times 10^{-3}}$

| **Cytokine** | **LPS vs. Control** | | **LPS+CBL vs. LPS** | | **LPS+CBL vs. Control** | |
|---|---|---|---|---|---|---|
| | **r** | ***p* value** | **r** | ***p* value** | **r** | ***p* value** |
| IL-10 | 0.70 | $\mathbf{1.4 \times 10^{-5}}$ ***** | 0.09 | $6.3 \times 10^{-1}$ | 0.70 | $\mathbf{1.5 \times 10^{-5}}$ ***** |
| IL-1α | 0.59 | $\mathbf{4.1 \times 10^{-4}}$ ***** | - 0.26 | $1.4 \times 10^{-1}$ | 0.46 | $7.4 \times 10^{-3}$ |
| IL-1β | 0.87 | $\mathbf{4.7 \times 10^{-10}}$ ***** | - 0.62 | $\mathbf{1.8 \times 10^{-4}}$ ***** | 0.87 | $\mathbf{7.0 \times 10^{-10}}$ ***** |
| IL-1RA | 0.84 | $\mathbf{9.5 \times 10^{-9}}$ ***** | - 0.60 | $\mathbf{3.1 \times 10^{-4}}$ ***** | 0.82 | $\mathbf{4.6 \times 10^{-8}}$ ***** |
| IL-4 | 0.66 | $\mathbf{4.5 \times 10^{-5}}$ ***** | 0.01 | $9.4 \times 10^{-1}$ | 0.59 | $\mathbf{3.3 \times 10^{-4}}$ ***** |
| IL-6 | 0.87 | $\mathbf{7.0 \times 10^{-10}}$ ***** | - 0.66 | $\mathbf{4.6 \times 10^{-5}}$ ***** | 0.87 | $\mathbf{7.0 \times 10^{-10}}$ ***** |
| IL-7 | 0.57 | $\mathbf{6.5 \times 10^{-4}}$ ***** | - 0.19 | $2.8 \times 10^{-1}$ | 0.67 | $\mathbf{2.9 \times 10^{-5}}$ ***** |
| IL-8 | 0.87 | $\mathbf{7.0 \times 10^{-10}}$ ***** | - 0.47 | $5.5 \times 10^{-3}$ | 0.86 | $\mathbf{2.3 \times 10^{-9}}$ ***** |
| IP-10 | 0.80 | $\mathbf{1.5 \times 10^{-7}}$ ***** | - 0.62 | $\mathbf{1.6 \times 10^{-4}}$ ***** | 0.63 | $\mathbf{1.3 \times 10^{-4}}$ ***** |
| MCP-1 | 0.70 | $\mathbf{1.1 \times 10^{-5}}$ ***** | - 0.55 | $\mathbf{1.1 \times 10^{-3}}$ ***** | 0.51 | $2.6 \times 10^{-3}$ |
| MIP-1α | 0.83 | $\mathbf{2.0 \times 10^{-8}}$ ***** | - 0.74 | $\mathbf{3.4 \times 10^{-6}}$ ***** | 0.83 | $\mathbf{3.2 \times 10^{-8}}$ ***** |
| MIP-1β | 0.87 | $\mathbf{2.3 \times 10^{-10}}$ ***** | - 0.78 | $\mathbf{3.6 \times 10^{-7}}$ ***** | 0.86 | $\mathbf{2.3 \times 10^{-9}}$ ***** |
| sCD40L | 0.10 | $5.6 \times 10^{-1}$ | 0.60 | $\mathbf{2.6 \times 10^{-4}}$ ***** | 0.40 | $2.2 \times 10^{-2}$ |
| TGF-α | 0.80 | $\mathbf{1.4 \times 10^{-7}}$ ***** | - 0.38 | $2.6 \times 10^{-2}$ | 0.71 | $\mathbf{7.0 \times 10^{-6}}$ ***** |
| TNF-α | 0.87 | $\mathbf{2.3 \times 10^{-10}}$ ***** | - 0.73 | $\mathbf{4.4 \times 10^{-6}}$ ***** | 0.86 | $\mathbf{1.2 \times 10^{-9}}$ ***** |

**Supplementary Table 5.** *Post hoc* pairwise comparisons of cytokine levels across 3 *ex vivo* conditions in healthy controls (n=33).

*****: significant at $\boldsymbol{\alpha} = \frac{\mathbf{0.05}}{\mathbf{15 \times 3}} \approx \mathbf{1.1} \times \mathbf{10^{-3}}$ (Bonferroni correction)

*r*: effect size computed from nonparametric test statistics

| Cytokine | LPS vs. Control | | LPS+CBL vs. LPS | | LPS+CBL vs. Control | |
|---|---|---|---|---|---|---|
| | r | *p* value | r | *p* value | r | *p* value |
| Fractalkine | 0.40 | **$7.0 \times 10^{-5}$ *** | - 0.21 | $4.4 \times 10^{-2}$ | 0.20 | $5.4 \times 10^{-2}$ |
| GM-CSF | 0.36 | **$3.6 \times 10^{-4}$ *** | - 0.15 | $1.5 \times 10^{-1}$ | 0.23 | $2.3 \times 10^{-2}$ |
| GRO | 0.46 | **$2.9 \times 10^{-6}$ *** | 0.16 | $1.2 \times 10^{-1}$ | 0.40 | **$7.1 \times 10^{-5}$ *** |
| IL-10 | 0.48 | **$1.0 \times 10^{-6}$ *** | - 0.06 | $5.3 \times 10^{-1}$ | 0.49 | **$5.3 \times 10^{-7}$ *** |
| IL-1α | 0.35 | **$5.2 \times 10^{-4}$ *** | - 0.21 | $3.9 \times 10^{-2}$ | 0.27 | $8.4 \times 10^{-3}$ |
| IL-1β | 0.80 | **$2.4 \times 10^{-20}$ **** | - 0.47 | **$2.0 \times 10^{-6}$ *** | 0.74 | **$1.5 \times 10^{-16}$ **** |
| IL-1RA | 0.69 | **$4.6 \times 10^{-14}$ **** | - 0.42 | **$1.8 \times 10^{-5}$ *** | 0.48 | **$1.0 \times 10^{-6}$ *** |
| IL-6 | 0.81 | **$1.2 \times 10^{-20}$ **** | - 0.61 | **$1.8 \times 10^{-10}$ *** | 0.73 | **$1.4 \times 10^{-15}$ **** |
| IL-7 | 0.36 | **$4.5 \times 10^{-4}$ *** | - 0.09 | $3.7 \times 10^{-1}$ | 0.27 | $9.6 \times 10^{-3}$ |
| IL-8 | 0.80 | **$5.0 \times 10^{-20}$ **** | - 0.19 | $5.6 \times 10^{-2}$ | 0.77 | **$1.1 \times 10^{-17}$ **** |
| IP-10 | 0.68 | **$2.6 \times 10^{-13}$ **** | - 0.53 | **$5.2 \times 10^{-8}$ *** | 0.27 | $8.1 \times 10^{-3}$ |
| MCP-1 | 0.72 | **$7.8 \times 10^{-15}$ **** | - 0.61 | **$2.5 \times 10^{-10}$ *** | 0.32 | $1.5 \times 10^{-3}$ |
| MIP-1α | 0.74 | **$2.6 \times 10^{-16}$ **** | - 0.63 | **$5.4 \times 10^{-11}$ **** | 0.58 | **$1.5 \times 10^{-9}$ *** |
| MIP-1β | 0.80 | **$3.2 \times 10^{-20}$ **** | - 0.71 | **$1.5 \times 10^{-14}$ **** | 0.68 | **$4.0 \times 10^{-13}$ **** |
| TGF-α | 0.60 | **$2.3 \times 10^{-10}$ *** | - 0.36 | **$3.5 \times 10^{-4}$ *** | 0.48 | **$8.3 \times 10^{-7}$ *** |
| TNF-α | 0.83 | **$8.8 \times 10^{-23}$ **** | - 0.69 | **$9.9 \times 10^{-14}$ **** | 0.75 | **$5.8 \times 10^{-17}$ **** |

**Supplementary Table 6.** *Post hoc* pairwise comparisons of cytokine levels across 3 *ex vivo* conditions in participants with psychiatric symptoms and/or diagnoses (n=97).
*: significant at $\boldsymbol{\alpha} = \frac{\mathbf{0.05}}{\mathbf{16 \times 3}} \approx \mathbf{1.0} \times \mathbf{10^{-3}}$ (Bonferroni correction)
**: significant at $\boldsymbol{\alpha} = \mathbf{1.0} \times \mathbf{10^{-10}}$
*r*: effect size computed from nonparametric test statistics

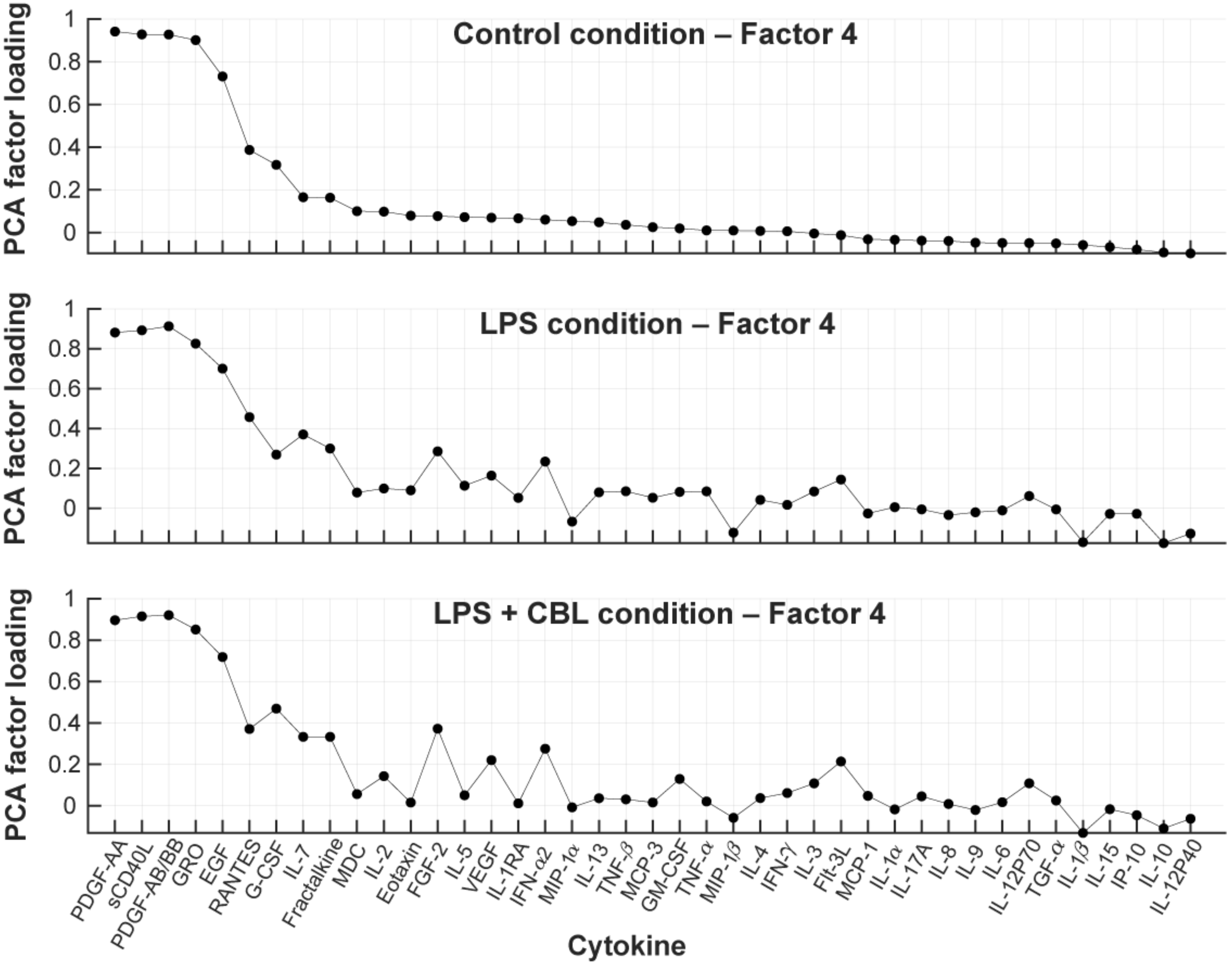

**Supplementary Figure 5**. Loadings of cytokine factors for which factor scores showed significant associations with anhedonia ratings after controlling for age, sex, and body mass index (BMI). For each condition, factor 4 was associated with at least one anhedonia scale. LPS+CBL Factor 4 was significantly associated with both anhedonia sub-components.

*Abbreviations*: CBL – clenbuterol; LPS – lipopolysaccharide; PCA – principal component analysis

| Control | | | | | LPS | | | | | LPS+CBL | | | | |
|---|---|---|---|---|---|---|---|---|---|---|---|---|---|---|
| **F1** | **F2** | **F3** | **F4** | **F5** | **F1** | **F2** | **F3** | **F4** | **F5** | **F1** | **F2** | **F3** | **F4** | **F5** |
| FGF-2 | Eotaxin | IL-1β | EGF | IL-1α | FGF-2 | Eotaxin | IL-1β | EGF | IL-1α | Eotaxin | GM-CSF | IL-1β | EGF | IL-1α |
| Flt-3L | IL-13 | IL-6 | GRO | IL-4 | Flt-3L | GM-CSF | IP-10 | G-CSF | IL-4 | FGF-2 | IL-13 | IP-10 | GRO | IL-4 |
| Fractalkine | IL-1RA | IP-10 | PDGF-AA | IL-9 | Fractalkine | IL-13 | MIP-1α | GRO | IL-9 | Flt-3L | IL-1RA | MIP-1α | PDGF-AA | IL-9 |
| G-CSF | IL-5 | MCP-1 | PDGF-AB/BB | | IFN-α2 | IL-1RA | MIP-1β | PDGF-AA | | Fractalkine | IL-5 | MIP-1β | PDGF-AB/BB | |
| GM-CSF | IL-8 | MIP-1α | RANTES | | IFN-γ | IL-5 | TNF-α | PDGF-AB/BB | | G-CSF | IL-6 | TNF-α | sCD40L | |
| IFN-α2 | MCP-3 | MIP-1β | sCD40L | | IL-10 | IL-6 | | RANTES | | IFN-α2 | IL-8 | | | |
| IFN-γ | TGF-α | TNF-α | | | IL-12P40 | IL-8 | | | | IFN-γ | MCP-1 | | | |
| IL-10 | TNF-β | | | | IL-12P70 | MCP-1 | | | | IL-10 | MCP-3 | | | |
| IL-12P40 | | | | | IL-15 | MCP-3 | | | | IL-12P40 | MDC | | | |
| IL-12P70 | | | | | IL-17A | MDC | | | | IL-12P70 | TGF-α | | | |
| IL-15 | | | | | IL-2 | TGF-α | | | | IL-15 | TNF-β | | | |
| IL-17A | | | | | IL-3 | TNF-β | | | | IL-17A | | | | |
| IL-2 | | | | | IL-7 | | | | | IL-2 | | | | |
| IL-3 | | | | | VEGF | | | | | IL-3 | | | | |
| IL-7 | | | | | | | | | | IL-7 | | | | |
| MDC | | | | | | | | | | RANTES | | | | |
| VEGF | | | | | | | | | | VEGF | | | | |

**Supplementary Table 7.** Major loadings of each cytokine factor in each condition. Factors with consistent loadings across conditions are marked with similar colors.

*Abbreviations*: EGF: epidermal growth factor; FGF: fibroblast growth factor; Flt3-L: FMS- like tyrosine kinase 3-ligand; G-CSF: Granulocyte colony-stimulating factor; GM-CSF: granulocyte-macrophage colony-stimulating factor; GRO: growth regulated oncogene; IFN: interferon; IL: interleukin; IP: interferon gamma-induced protein; MCP: monocyte chemotactic protein; MDC: macrophage-derived chemokine; MIP: macrophage inflammatory protein; PDGF: platelet-derived growth factor; RANTES: regulated on activation, normal T cell expressed and secreted; sCD40L: soluble cluster of differentiation 40 ligand; TGF: transforming growth factor; TNF: tumor necrosis factor; VEGF: vascular endothelial growth factor.

*For ease of interpretation, all factors in LPS and LPS+CBL conditions were listed following the same numbering scheme as Control factors.

| Factors | Control | | LPS | | LPS + CBL | |
|---|---|---|---|---|---|---|
| | *rho* | *p* value | *rho* | *p* value | *rho* | *p* value |
| **Controlling for Age, Sex, and BMI** | | | | | | |
| **BDI (Depression)** | | | | | | |
| F1 | - 0.10 | 0.26 | - 0.11 | 0.24 | - 0.14 | 0.12 |
| F2 | - 0.064 | 0.49 | $- 3.2 \times 10^{-3}$ | 0.97 | 0.029 | 0.75 |
| F3 | 0.028 | 0.76 | - 0.020 | 0.83 | - 0.023 | 0.80 |
| F4 | 0.14 | 0.12 | 0.16 | 0.088 | 0.17 | 0.067 |
| F5 | 0.14 | 0.12 | 0.032 | 0.72 | 0.020 | 0.83 |
| **MASC (Anxiety)** | | | | | | |
| F1 | - 0.22 | 0.015 | - 0.20 | 0.031 | - 0.27 | $3.7 \times 10^{-3}$ |
| F2 | - 0.012 | 0.90 | 0.13 | 0.17 | 0.17 | 0.062 |
| F3 | - 0.022 | 0.82 | - 0.10 | 0.29 | - 0.14 | 0.13 |
| F4 | 0.033 | 0.73 | 0.078 | 0.41 | 0.067 | 0.48 |
| F5 | 0.096 | 0.30 | 0.015 | 0.87 | - 0.026 | 0.78 |
| **TEPS-A (Anticipatory Anhedonia)** | | | | | | |
| F1 | 0.044 | 0.66 | 0.046 | 0.66 | 0.063 | 0.54 |
| F2 | 0.035 | 0.74 | - 0.019 | 0.85 | - 0.068 | 0.50 |
| F3 | 0.031 | 0.76 | 0.11 | 0.27 | 0.13 | 0.21 |
| F4 | **- 0.35** | **$3.6 \times 10^{-4}$ *** | **- 0.36** | **$2.6 \times 10^{-4}$ *** | **- 0.39** | **$7.4 \times 10^{-5}$ *** |
| F5 | -0.16 | 0.12 | $7.0 \times 10^{-3}$ | 0.95 | - 0.057 | 0.57 |
| **TEPS-C (Consummatory Anhedonia)** | | | | | | |
| F1 | - 0.11 | 0.31 | - 0.067 | 0.52 | - 0.028 | 0.79 |
| F2 | 0.23 | 0.024 | 0.11 | 0.32 | 0.079 | 0.44 |
| F3 | 0.18 | 0.081 | 0.016 | 0.88 | - 0.050 | 0.63 |
| F4 | - 0.29 | $4.3 \times 10^{-3}$ | - 0.30 | $3.9 \times 10^{-3}$ | **- 0.36** | **$3.3 \times 10^{-4}$ *** |
| F5 | - 0.19 | 0.058 | - 0.020 | 0.85 | - 0.15 | 0.14 |
| **Controlling for Age, Sex, BMI, and Depression Severity** | | | | | | |
| **TEPS-A** | | | | | | |
| F1 | 0.049 | 0.63 | 0.043 | 0.68 | 0.033 | 0.75 |
| F2 | $- 4.7 \times 10^{-3}$ | 0.96 | - 0.043 | 0.68 | - 0.078 | 0.45 |
| F3 | 0.070 | 0.50 | 0.11 | 0.29 | 0.13 | 0.22 |
| F4 | - 0.31 | $1.7 \times 10^{-3}$ | - 0.32 | $1.8 \times 10^{-3}$ | **- 0.34** | **$6.6 \times 10^{-4}$ #** |
| F5 | - 0.064 | 0.54 | 0.068 | 0.51 | 0.016 | 0.88 |
| **TEPS-C** | | | | | | |
| F1 | - 0.12 | 0.26 | - 0.091 | 0.39 | - 0.052 | 0.62 |
| F2 | 0.23 | 0.024 | 0.12 | 0.27 | 0.082 | 0.43 |
| F3 | 0.19 | 0.066 | $- 3.4 \times 10^{-3}$ | 0.98 | - 0.086 | 0.41 |
| F4 | - 0.27 | $9.2 \times 10^{-3}$ | - 0.26 | 0.012 | **- 0.34** | **$9.1 \times 10^{-4}$ #** |
| F5 | - 0.16 | 0.13 | $1.0 \times 10^{-3}$ | 0.99 | - 0.14 | 0.20 |

**Supplementary Table 8.** Spearman correlations between cytokine factors of the three conditions and a) depression, b) anxiety, c) anticipatory anhedonia, and d) consummatory anhedonia.

* significant at $\alpha = \frac{0.05}{5\times4\times3} \approx \mathbf{8.3} \times \mathbf{10^{-4}}$

### significant at $\alpha = \frac{0.05}{5\times2\times3} \approx \mathbf{1.7} \times \mathbf{10^{-3}}$

*Abbreviations*: BDI: Beck Depression Inventory; BMI: Body Mass Index (kg/m²); MASC: Multidimensional Anxiety Scale for Children; TEPS-A: Temporal Experience of Pleasure Scale – Anticipatory; TEPS-C: Temporal Experience of Pleasure Scale – Consummatory

| Factors | $\frac{\textbf{LPS}}{\textbf{LPS + CBL}}$ | |
|---|---|---|
| | *rho* | *p* value |
| **Controlling for Age, Sex, and BMI** | | |
| **TEPS-A (Anticipatory Anhedonia)** | | |
| F1 | -0.04 | 0.73 |
| F2 | 0.02 | 0.85 |
| F3 | -0.17 | 0.12 |
| F4 | -0.04 | 0.74 |
| **TEPS-C (Consummatory Anhedonia)** | | |
| F1 | -0.07 | 0.55 |
| F2 | -0.19 | 0.08 |
| F3 | -0.12 | 0.26 |
| F4 | 0.08 | 0.45 |
| **Controlling for Age, Sex, BMI, and Depression Severity** | | |
| **TEPS-A (Anticipatory Anhedonia)** | | |
| F1 | -0.04 | 0.69 |
| F2 | 0.11 | 0.28 |
| F3 | **-0.26** | **0.014** |
| F4 | -0.09 | 0.51 |
| **TEPS-C (Consummatory Anhedonia)** | | |
| F1 | -0.04 | 0.70 |
| F2 | -0.11 | 0.30 |
| F3 | -0.15 | 0.16 |
| F4 | 0.08 | 0.42 |

**Supplementary Table 9.** Spearman correlations between anhedonia scores and 4 immune factors derived from the ratio of cytokine levels between LPS and LPS+CBL conditions. At the uncorrected level of $p < 0.05$, Ratio F3 (bolded) showed significant association with anticipatory anhedonia adjusting for age, sex, BMI, and depression. There is no significant association at the Bonferroni-corrected significant level, $\alpha = \frac{\mathbf{0.05}}{\mathbf{4}} = \mathbf{0.0125}$.
*Abbreviations*: BDI: Beck Depression Inventory; BMI: Body Mass Index (kg/m²); MASC: Multidimensional Anxiety Scale for Children; TEPS-A: Temporal Experience of Pleasure Scale – Anticipatory; TEPS-C: Temporal Experience of Pleasure Scale – Consummatory

| | | | | | P-values | | | | | |
|---|---|---|---|---|---|---|---|---|---|---|
| *Δrho* | F2 – F1 | F3 – F1 | F3 – F2 | F4 – F1 | F4 – F2 | F4 – F3 | F5 – F1 | F5 – F2 | F5 – F3 | F5 – F4 |
| **Control** | | | | | | | | | | |
| Controlling for Age, Sex, and BMI | | | | | | | | | | |
| BDI | 0.79 | 0.23 | 0.56 | 0.04 | 0.17 | 0.37 | 0.05 | 0.17 | 0.42 | 0.99 |
| MASC | 0.18 | 0.07 | 0.95 | 0.05 | 0.77 | 0.67 | $6.11 \times 10^{-3}$ | 0.51 | 0.40 | 0.63 |
| TEPS-A | 0.97 | 0.93 | 0.97 | $\mathbf{2.09 \times 10^{-3}}$* | 0.02 | $6.32 \times 10^{-3}$ | 0.11 | 0.30 | 0.22 | 0.03 |
| TEPS-C | 0.06 | 0.02 | 0.73 | 0.22 | $\mathbf{4.75 \times 10^{-4}}$* | $\mathbf{1.04 \times 10^{-3}}$* | 0.48 | 0.02 | 0.01 | 0.35 |
| Controlling for Age, Sex, BMI, and Depression Severity | | | | | | | | | | |
| TEPS-A | 0.77 | 0.88 | 0.64 | $6.63 \times 10^{-3}$ | 0.06 | $5.68 \times 10^{-3}$ | 0.33 | 0.73 | 0.38 | 0.01 |
| TEPS-C | 0.08 | 0.02 | 0.79 | 0.36 | $\mathbf{4.48 \times 10^{-4}}$* | $\mathbf{2.38 \times 10^{-3}}$* | 0.74 | 0.03 | 0.03 | 0.33 |
| **LPS** | | | | | | | | | | |
| Controlling for Age, Sex, and BMI | | | | | | | | | | |
| BDI | 0.55 | 0.50 | 0.92 | 0.06 | 0.27 | 0.17 | 0.30 | 0.78 | 0.70 | 0.36 |
| MASC | 0.05 | 0.39 | 0.15 | 0.05 | 0.74 | 0.19 | 0.08 | 0.41 | 0.36 | 0.66 |
| TEPS-A | 0.73 | 0.63 | 0.48 | $8.30 \times 10^{-3}$ | 0.02 | $\mathbf{4.83 \times 10^{-4}}$* | 0.78 | 0.88 | 0.40 | $\mathbf{2.97 \times 10^{-4}}$* |
| TEPS-C | 0.40 | 0.59 | 0.58 | 0.16 | $\mathbf{1.53 \times 10^{-3}}$* | 0.02 | 0.77 | 0.41 | 0.77 | $\mathbf{4.95 \times 10^{-3}}$* |
| Controlling for Age, Sex, BMI, and Depression Severity | | | | | | | | | | |
| TEPS-A | 0.66 | 0.65 | 0.41 | 0.03 | 0.06 | $\mathbf{2.31 \times 10^{-3}}$* | 0.86 | 0.52 | 0.70 | $\mathbf{1.46 \times 10^{-3}}$* |
| TEPS-C | 0.35 | 0.61 | 0.47 | 0.32 | $\mathbf{2.61 \times 10^{-3}}$* | 0.04 | 0.56 | 0.47 | 0.99 | 0.01 |
| **LPS+CBL** | | | | | | | | | | |
| Controlling for Age, Sex, and BMI | | | | | | | | | | |
| BDI | 0.31 | 0.36 | 0.75 | 0.02 | 0.34 | 0.15 | 0.21 | 0.94 | 0.76 | 0.21 |
| MASC | $7.61 \times 10^{-3}$ | 0.28 | 0.04 | 0.02 | 0.43 | 0.13 | 0.05 | 0.14 | 0.37 | 0.45 |
| TEPS-A | 0.46 | 0.66 | 0.28 | $\mathbf{2.22 \times 10^{-3}}$* | 0.03 | $\mathbf{2.10 \times 10^{-4}}$* | 0.41 | 0.95 | 0.17 | $\mathbf{6.54 \times 10^{-4}}$* |
| TEPS-C | 0.56 | 0.89 | 0.45 | 0.04 | $\mathbf{1.05 \times 10^{-3}}$* | 0.04 | 0.39 | 0.14 | 0.49 | 0.03 |
| Controlling for Age, Sex, BMI, and Depression Severity | | | | | | | | | | |
| TEPS-A | 0.55 | 0.57 | 0.22 | 0.01 | 0.08 | $\mathbf{1.46 \times 10^{-3}}$* | 0.90 | 0.53 | 0.42 | $\mathbf{2.08 \times 10^{-3}}$* |
| TEPS-C | 0.53 | 0.83 | 0.30 | 0.08 | $\mathbf{3.07 \times 10^{-3}}$* | 0.08 | 0.57 | 0.19 | 0.74 | 0.06 |

**Supplementary Table 10**. *P*-values of differences (*Δrho*) between factor-symptom associations across conditions. *P*-values were generated from *post-hoc* bias-corrected and accelerated bootstrapping method with $10^5$ resamples. Factor 4 was determined to have significant associations with anhedonia in original analyses.

* Significant at $\alpha = \frac{0.05}{10} = \mathbf{5.0 \times 10^{-3}}$

*Abbreviations*: BDI: Beck Depression Inventory; BMI: Body Mass Index (kg/m²); MASC: Multidimensional Anxiety Scale for Children; TEPS-A: Temporal Experience of Pleasure Scale – Anticipatory; TEPS-C: Temporal Experience of Pleasure Scale – Consummatory

| | F1 | | | F2 | | | F3 | | | F4 | | | F5 | | |
|---|---|---|---|---|---|---|---|---|---|---|---|---|---|---|---|
| | | 95% CI | | | 95% CI | | | 95% CI | | | 95% CI | | | 95% CI | |
| | *rho* | Lower Limit | Upper Limit | *rho* | Lower Limit | Upper Limit | *rho* | Lower Limit | Upper Limit | *rho* | Lower Limit | Upper Limit | *rho* | Lower Limit | Upper Limit |
| **Control** | | | | | | | | | | | | | | | |
| Controlling for Age, Sex, and BMI | | | | | | | | | | | | | | | |
| BDI | - 0.10 | -0.28 | 0.08 | - 0.06 | -0.26 | 0.13 | 0.03 | -0.17 | 0.22 | 0.14 | -0.04 | 0.31 | 0.14 | -0.04 | 0.31 |
| MASC | - 0.22 | -0.39 | -0.05 | - 0.01 | -0.21 | 0.19 | - 0.02 | -0.20 | 0.16 | 0.03 | -0.16 | 0.23 | 0.10 | -0.10 | 0.28 |
| TEPS-A | 0.04 | -0.15 | 0.22 | 0.04 | -0.18 | 0.26 | 0.031 | -0.18 | 0.23 | **- 0.35** | -0.52 | -0.16 | -0.16 | -0.35 | 0.06 |
| TEPS-C | - 0.11 | -0.31 | 0.11 | 0.23 | 0.018 | 0.43 | 0.18 | -0.01 | 0.36 | - 0.29 | -0.47 | -0.09 | - 0.19 | -0.39 | 0.01 |
| Controlling for Age, Sex, BMI, and Depression Severity | | | | | | | | | | | | | | | |
| TEPS-A | 0.05 | -0.15 | 0.24 | - 4.7 × $10^{-3}$ | -0.22 | 0.22 | 0.07 | -0.14 | 0.27 | - 0.31 | -0.50 | -0.11 | - 0.06 | -0.27 | 0.14 |
| TEPS-C | - 0.12 | -0.33 | 0.13 | 0.23 | 0.02 | 0.43 | 0.19 | -0.01 | 0.38 | - 0.27 | -0.45 | -0.07 | - 0.16 | -0.35 | 0.04 |
| **LPS** | | | | | | | | | | | | | | | |
| Controlling for Age, Sex, and BMI | | | | | | | | | | | | | | | |
| BDI | - 0.11 | -0.29 | 0.08 | - 3.2 × $10^{-3}$ | -0.20 | 0.20 | - 0.02 | -0.21 | 0.17 | 0.16 | -0.03 | 0.34 | 0.03 | -0.16 | 0.22 |
| MASC | - 0.20 | -0.37 | -0.02 | 0.13 | -0.06 | 0.31 | - 0.10 | -0.29 | 0.10 | 0.08 | -0.13 | 0.28 | 0.02 | -0.18 | 0.20 |
| TEPS-A | 0.05 | -0.15 | 0.24 | - 0.02 | -0.24 | 0.21 | 0.11 | -0.11 | 0.33 | **- 0.36** | -0.53 | -0.16 | 7.0 × $10^{-3}$ | -0.20 | 0.21 |
| TEPS-C | - 0.07 | -0.27 | 0.15 | 0.11 | -0.11 | 0.31 | 0.02 | -0.20 | 0.23 | - 0.30 | -0.47 | -0.11 | - 0.020 | -0.21 | 0.17 |
| Controlling for Age, Sex, BMI, and Depression Severity | | | | | | | | | | | | | | | |
| TEPS-A | 0.04 | -0.16 | 0.25 | - 0.04 | -0.29 | 0.20 | 0.11 | -0.12 | 0.33 | - 0.32 | -0.51 | -0.10 | 0.07 | -0.15 | 0.27 |
| TEPS-C | - 0.09 | -0.30 | 0.15 | 0.12 | -0.11 | 0.33 | - 3.4 × $10^{-3}$ | -0.21 | 0.21 | - 0.26 | -0.44 | -0.07 | 1.0 × $10^{-3}$ | -0.19 | 0.19 |
| **LPS+CBL** | | | | | | | | | | | | | | | |
| Controlling for Age, Sex, and BMI | | | | | | | | | | | | | | | |
| BDI | -0.14 | -0.31 | 0.04 | 0.03 | -0.16 | 0.22 | - 0.02 | -0.22 | 0.17 | 0.17 | -0.02 | 0.34 | 0.02 | -0.17 | 0.21 |
| MASC | - 0.27 | -0.43 | -0.09 | 0.17 | -0.01 | 0.34 | - 0.14 | -0.32 | 0.05 | 0.07 | -0.13 | 0.26 | - 0.03 | -0.21 | 0.16 |
| TEPS-A | 0.06 | -0.13 | 0.25 | - 0.07 | -0.27 | 0.15 | 0.13 | -0.10 | 0.34 | **- 0.39** | -0.55 | -0.19 | - 0.06 | -0.25 | 0.14 |
| TEPS-C | - 0.03 | -0.23 | 0.19 | 0.08 | -0.13 | 0.28 | - 0.05 | -0.27 | 0.18 | **- 0.36** | -0.53 | -0.16 | - 0.15 | -0.35 | 0.05 |
| Controlling for Age, Sex, BMI, and Depression Severity | | | | | | | | | | | | | | | |
| TEPS-A | 0.03 | -0.16 | 0.24 | - 0.08 | -0.30 | 0.15 | 0.13 | -0.09 | 0.34 | **- 0.34** | -0.52 | -0.13 | 0.02 | -0.18 | 0.21 |
| TEPS-C | - 0.05 | -0.26 | 0.18 | 0.08 | -0.14 | 0.29 | - 0.09 | -0.29 | 0.14 | **- 0.34** | -0.50 | -0.14 | - 0.14 | -0.33 | 0.07 |

**Supplementary Table 11**. Factor-symptom correlation coefficients across conditions and their corresponding 95% confidence intervals, computed using *post-hoc* bias-corrected and accelerated bootstrap ($10^5$ resamples). Bolded *rho* values denote significant associations in original analyses. *Abbreviations*: BDI: Beck Depression Inventory; BMI: Body Mass Index (kg/m²); MASC: Multidimensional Anxiety Scale for Children; TEPS-A: Temporal Experience of Pleasure Scale – Anticipatory; TEPS-C: Temporal Experience of Pleasure Scale – Consummatory
